# Disease-specific selective vulnerability and neuroimmune pathways in dementia revealed by single cell genomics

**DOI:** 10.1101/2023.09.29.560245

**Authors:** Jessica E. Rexach, Yuyan Cheng, Lawrence Chen, Damon Polioudakis, Li-Chun Lin, Vivianne Mitri, Andrew Elkins, Anna Yin, Daniela Calini, Riki Kawaguchi, Jing Ou, Jerry Huang, Christopher Williams, John Robinson, Stephanie E. Gaus, Salvatore Spina, Edward B. Lee, Lea T. Grinberg, Harry Vinters, John Q. Trojanowski, William W. Seeley, Dheeraj Malhotra, Daniel H. Geschwind

## Abstract

The development of successful therapeutics for dementias requires an understanding of their shared and distinct molecular features in the human brain. We performed single-nuclear RNAseq and ATACseq in Alzheimer disease (AD), Frontotemporal degeneration (FTD), and Progressive Supranuclear Palsy (PSP), analyzing 40 participants, yielding over 1.4M cells from three brain regions ranging in vulnerability and pathological burden. We identify 35 shared disease-associated cell types and 14 that are disease-specific, replicating those previously identified in AD. Disease_-_specific cell states represent molecular features of disease-specific glial-immune mechanisms and neuronal vulnerability in each disorder, layer 4/5 intra-telencephalic neurons in AD, layer 2/3 intra-telencephalic neurons in FTD, and layer 5/6 near-projection neurons in PSP. We infer intrinsic disease-associated gene regulatory networks, which we empirically validate by chromatin footprinting. We find that causal genetic risk acts in specific neuronal and glial cells that differ across disorders, primarily non-neuronal cells in AD and specific neuronal subtypes in FTD and PSP. These data illustrate the heterogeneous spectrum of glial and neuronal composition and gene expression alterations in different dementias and identify new therapeutic targets by revealing shared and disease-specific cell states.

## Introduction

Alzheimer Disease (AD), Frontotemporal Dementia (FTD) and progressive supranuclear palsy (PSP) are clinical syndromes involving distinct neuropathologically defined conditions that involve different forms of tau pathology (Chung et al., 2021). AD and PSP are canonically defined by tau pathology, whereas FTD may display either TDP (FTLD-TDP) or Tau (FTLD-Tau) pathological inclusions in post-mortem brain, the latter of which is typically observed in case with the behavioral variant of FTD (bvFTD) (Bahia et al., 2013). Combined, these three disorders affect over 28 million people worldwide (Young et al., 2018). Compared to AD where 143 drugs are in active clinical trials in 2022, PSP and FTD have only a handful of drugs in development (Boeve et al., 2022). Contributing to the challenge of therapeutic development is a limited molecular understanding of FTD and PSP, coupled to not knowing what neuroinflammatory or neurodegenerative features are similar or distinct in in each disorder.

A foundational observation in dementia is the presence of selective neuronal vulnerability, wherein neurodegeneration, tau pathology and neuroinflammation impact specific cell types and brain regions in temporally and spatially distinct patterns in each disorder, leading to distinct clinical syndromes (Fu et al., 2018; Rexach and Geschwind, 2020). In addition, both AD and bvFTD have distinct patterns of cortical layer-specific pathology (Santillo and Englund, 2014), superficial layer cortical projection neurons being more vulnerable in bvFTD (Kersaitis et al., 2004) whereas in PSP, frontal cortical and subcortical motor pathways are more vulnerable (Chung et al., 2021; Kovacs et al., 2020). In bvFTD, the insular cortex and salience network are differentially vulnerable (Zhou et al., 2010). Furthermore, the genetic architectures of AD, PSP and bvFTD are distinct, highlighting differential contributions of neuronal and glia cell types to disease risk (Cooper et al., 2022; Endo et al., 2022; Huang et al., 2017; Rexach et al., 2020; Swarup et al., 2020; Swarup et al., 2019). Despite these differences, all three disorders involve tau pathology that begins in more vulnerable areas and then progresses, potentially through shared prion-like spreading phenomena and/or additional unknown mechanisms (Chung *et al*., 2021; Kaufman et al., 2016; Kim et al., 2020; Kovacs *et al*., 2020).

Advances in single-cell sequencing have revealed initial candidate markers of selective vulnerability in AD, including regulators of neuronal transcription (Leng et al., 2021; Morabito et al., 2021), gene signatures of neurons containing hyperphosphorylated tau (Otero-Garcia et al., 2022) and sex-specific glial vulnerability patterns (Mathys et al., 2019). Neural-immune activation is another universal early and persistent feature in dementia and heterogenous glial types have been described in post-mortem AD brain (Grubman et al., 2019; Mathys *et al*., 2019; Morabito *et al*., 2021; Nguyen et al., 2020; Olah et al., 2020; Patrick et al., 2021; Sadick et al., 2022; Zhou et al., 2020). Despite these advances (Grubman *et al*., 2019; Leng *et al*., 2021; Mathys *et al*., 2019; Otero-Garcia *et al*., 2022; Zhou *et al*., 2020), studies have yet to formally compare across disorders involving tau. Thus, much remains unknown, including the specificity of changes observed in AD compared with other disorders, their role in selective neuronal vulnerability, or glial diversity. We address these important gaps in knowledge through direct comparison of post-mortem brain at the single cell level across three major disorders involving tau pathology, AD, bvFTD and PSP, using both snRNAseq and snATACseq to validate shared and specific disease-associated cell states and their predicted regulatory drivers. Our design enables identification of new shared and distinct markers of neuronal vulnerability, glial states that vary across disease and disorder-specific cellular differences in the expression and regulation of known risk genes.

## Results

We compared three disorders that are collectively referred to as tauopathies, but that display different regional and laminar patterns of neuronal loss and glial activation (Figure 1A): AD, bvFTD and PSP (Chung *et al*., 2021; Leng *et al*., 2021) using single nuclear sequencing of mRNA (snSeq) in conjunction with the Assay for Transposase-Accessible Chromatin (ATACseq; Buenrostro et al., 2015; Stuart et al., 2019). Using a well curated, neuropathologically-characterized brain collection (Methods), we selected three cortical brain regions with differential vulnerability to disease (Braak et al., 2006; Kim *et al*., 2020; Kovacs *et al*., 2020) and prospective, semi-quantitative ratings of burden of tau protein hyperphosphorylation (tau score) and microvacuolations, astrogliosis, and neuronal loss (neurodegeneration score) per sample using a published grading scale ((Kim *et al*., 2020; Lin et al., 2019); see Methods). The distribution of pathology scores matched the expected distribution of pathology in each disorder based on published annotations (Figure 1B, Table S1). For example, the insular cortex showed the highest disease burden in bvFTD (Kim *et al*., 2020) and moderate burden in PSP and AD. In all three disorders, the motor cortex (primary motor cortex (M1; BA4) had comparable, moderate levels of tau pathology, which in PSP was the highest of the three cortical regions profiled, as expected (Kovacs *et al*., 2020). In contrast, the primary visual cortex (V1), a relatively spared region, had low levels of tau pathology in all three disorders, and we hypothesized that it would have higher expression of cellular resilience factors compared to the more vulnerable brain regions.

**Figure 1:**
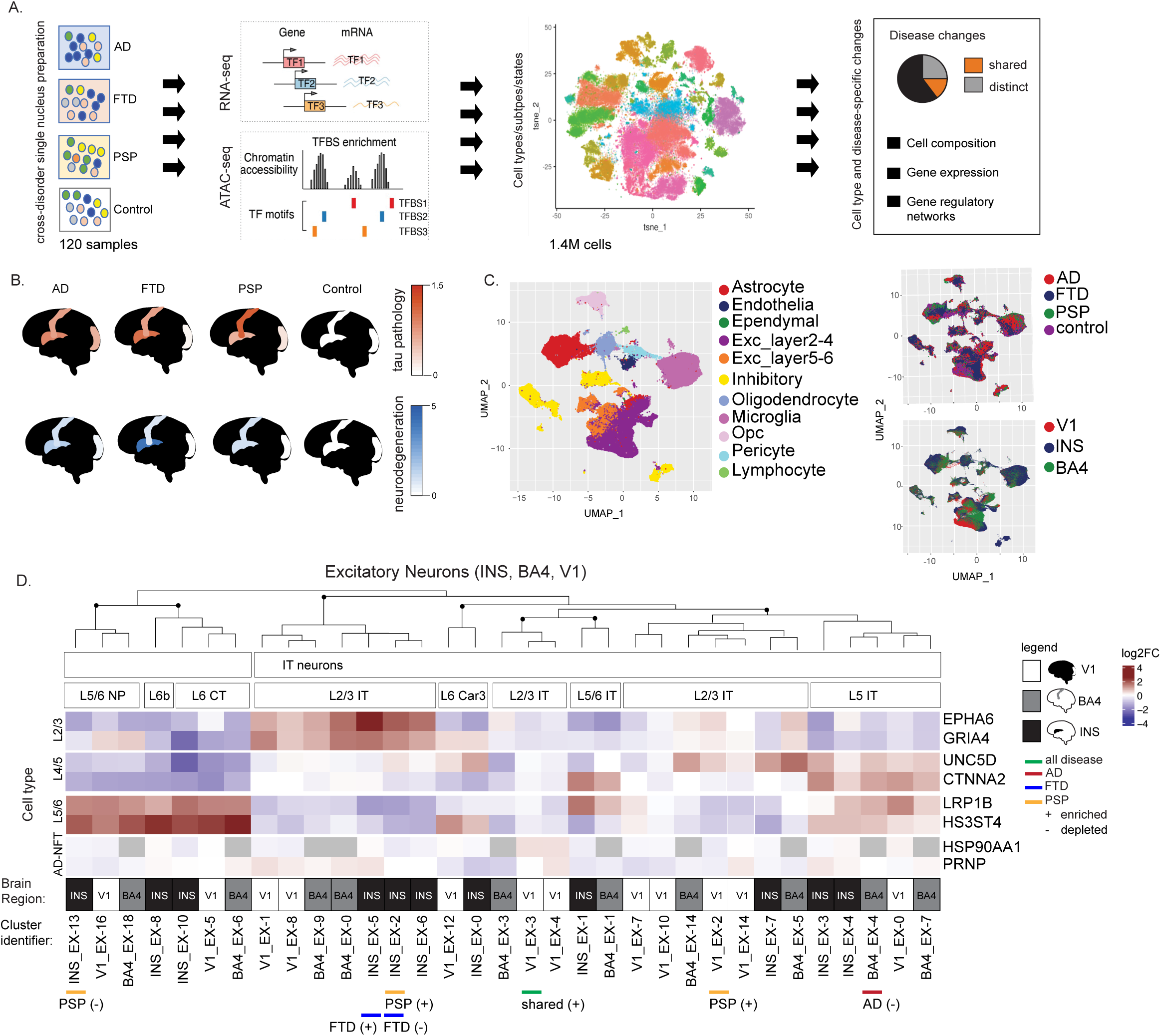
Comparison of cell types, subclasses and disease states across brain regions with variable disease vulnerability across neurodegenerative tauopathies. **(A)** Schema depicting cross-disorder multiomic analysis of post-mortem human brain tissue from three neurodegenerative tauopathies (AD, bvFTD, PSP, controls; 120 samples, Table S1) including three cortical brain regions (INS, BA4, V1) using single nuclear RNA-seq (snRNA-seq) and ATAC-seq (1.4M cells) to define and distinguish disorder-specific and shared changes in cellular molecular composition, gene expression and gene regulatory networks. **(B)** Cartoon with heatmap showing average neuropathology scores for neurodegeneration (blue, below) and tau (red above) measured across subjects, by region and disorder (Methods; (Table S1). **(C)** UMAP of snRNA-seq clusters separating into 11 major cell types (colored per legend; Methods) and including nuclei from each disease condition (AD, bvFTD, PSP, control; right top) and brain region (V1, INS, BA4; right lower). **(D)** Hierarchical clustering of excitatory neuron clusters from different brain regions (INS in black; BA4 in grey, and V1 in white; as shown in key) into reference-based subclasses and related clusters, showing 8 example marker genes variable expressed by groups including markers separating neuronal layers (L2/3; L4/5, L5/6) and markers of neurons bearing tau neurofibrillary tangles (AD_NFT) defined in AD brain (Otero-Garcia *et al*., 2022) (Methods, Figure S1C-E for cluster assignment to reference human motor cortex cell subclasses (Bakken *et al*., 2021); IT = intratelencephalic NP = near projecting, CT = corticothalamic, L2/3, L4/5, L5/6 refers to layer-specific neuronal classes; extratelencephalic (ET) neurons contribute cells to BA4_EX3 and BA4_EX-1 as shown in Figure S2E but do not populate a distinct cluster). Clusters significantly depleted (-) or enriched (+) in disease cells are indicated with a colored line (AD red; bvFTD blue, PSP gold; (limma, FDR <0.1, see Table S3 for complete differential cluster composition results; bvFTD abbreviated to “FTD” throughout figures).

We generated 880,000 single nuclear mRNA expression profiles from 120 brain samples representing 40 subjects (10 from each disorder and 10 control subjects), with three cortical regions from each (Figure 1, Figure S1, Methods). Following stringent quality control and outlier removal (Methods), 590,541 high quality cell profiles from 107 patient samples remained (Table S1). Confounding variables of age, sex, postmortem interval and RIN were not significantly different between control and subjects (*p* < 0.05, Wilcoxon, Table S1).We used human primary motor cortex for reference-based single cell analysis (Methods, Figure S1C,D,E, Figure 1D, Table S2) because it has been deeply profiled as part of the BRAIN initiative to establish conserved clusters with standardized nomenclature (Bakken et al., 2021; Network, 2021). We identified 9 canonical cell classes, including both abundant and rare cell subtypes (excitatory neurons (EX), inhibitory neurons (IN), astrocytes (AST), oligodendrocytes (OL), oligodendrocyte progenitor cells (OPC), microglia (MIC), endothelial cells (END), pericytes and lymphocytes (Hodge et al., 2019; Kelley et al., 2018; Mathys et al., 2019; Sweeney et al., 2016) and 24 canonical cell subclasses (Figure 1C, Figure S1C), which overlap with established subclasses from human brain (Bakken et al., 2021; Network, 2021) (Figure S1E).

### Molecular taxonomy of CNS cell types across brain regions in dementia

With robust identification of all major CNS cell types and subclasses, we leveraged our multi-region, multi-disease design to identify both shared and novel disease and brain region associated cell states. We reasoned that integration and re-clustering of each of the 9 major cell types independently in each of the three brain regions would maximize the likelihood of finding novel disease associated cell-states. We identified 181 clusters among 9 major cell types from 4 conditions (3 disorders and controls, excluding lymphocytes for low abundance), and 3 brain regions (Methods). The 181 clusters include 33 excitatory neurons, 26 inhibitory neurons, 56 oligodendroglia, 28 astrocytes, 18 microglia, 13 endothelial and 5 pericyte clusters. We then performed hierarchical clustering within major cell classes and used marker genes to group clusters from different brain regions to define an unbiased nomenclature based on the condition, region, cell type and cluster number that distinguishes each cluster. For example, an excitatory neuronal cluster from BA4 is labeled BA4_EX-numeric, ending with a unique numeric identifier that was assigned during clustering based on relative cluster size (BA4/V1/INS-EX/IN/OL/AST/OPC/MIC – 0-15 (Figure 1, Figure S2; Methods). As expected, the first order of clustering was driven predominantly by cell subclass, such as parvalbumin interneuron (Pvalb), or protoplasmic astrocyte (Figure 1D, Figure S2A-H, Table S2). At the next branch, clusters were divided further by brain region, with the majority of clusters identified in multiple brain regions (94%), with only 3 states restricted to only one region, one representing AST, one IN, and one MIC (Figure S2).

The 33 excitatory and 26 inhibitory neuronal clusters represent all canonical neuronal subclasses (Figure 1D, Figure S1A, Table S2, (Bakken *et al*., 2021)). 56 oligodendroglia clusters all belong to the three known subtypes of OL i.e early myelinating (Fard et al., 2017) *BCAS1*+ OL, as seen adjacent to chronic MS plaques (Figure S2B), and more mature OL expressing either high *PLP1* or *RBFOX1*, separated into 37 clusters based on brain region and state. We identify 2 subclasses of immature and differentiating OPC, clustered into 17 state- and region-dependent groups based on their distinct expression of known markers of proliferation, NMDA-directed migration (Xiao et al., 2013), neuroprotection (Rupnik et al., 2021) and axon interaction and myelination (Huang et al., 2020), (Figure S1D). The 28 astrocyte clusters are divided across 2 canonical subtypes, protoplasmic and fibrous astrocytes. Protoplasmic astrocytes exist in 4 transcriptomically-distinct groups suggestive of functionally variable states, and fibrous astrocytes in 1 group (Figure S2C, Table S2). The 18 microglial clusters are grouped into six transcriptomically distinct states (Figure S1G; methods; Table S2). Reassuringly, each of these six transcriptomically-distinct microglial states overlapped significantly with those previously profiled from fresh human brain tissue (Olah et al., 2020), supporting the quality of our data (Table S4). Among these, five overlapped significantly with microglia clusters previously identified in frontal cortex in AD (Mathys et al., 2019) (Table S4), including a distinct cluster marked by neuropathologically validated “motile” microglia markers *NEAT1*/*FDG4* compared to the “dystrophic” microglia marker *FLT* (Nguyen et al., 2020). The 13 clusters of endothelial cells are divided into two major groups marked by either higher expression of immune signaling genes or genes involved in angiogenesis and endothelial maintenance (Figure S2E) (Fan et al., 2014; Zhao et al., 2015) and 5 clusters of pericytes (Figure S2F; Table S2). In summary, we performed a systematic classification and annotation of 181 cell states representing all established neuronal and glial cell types and subclasses, their regional localization in three human brain regions, their distinguishing marker genes, associated biological pathways, transcriptional regulators, and differential gene expression (DGE) across disorders (Table S2, S3, S5).

We first identified genes that were significantly differentially expressed (DE) in major cell types across subjects by disorder, using a stringent model to correct for multiple comparisons (Methods). It was notable that disorder-specific differentially expressed genes were a minority; the vast majority of DE genes were shared by more than one disorder (95% of 6,081 genes; FDR <0.05 and Log2FC>0.1, Cross-Disorder LME, Table S5), indicating that these tauopathies have substantial shared molecular pathology. Of the 5% of genes detected as DE in only one of the disorders, the majority (92%; 280 genes) were DE in only a single cell type (Table S5). Notable examples of such disorder specific DE genes in microglia include *PTPRG* and *IL15* in AD (BA4-microglia, INS-microglia; respectively), consistent with recent observations (Wendimu and Hooks, 2022). Other genes DE in microglia in a disease-specific manner include *VPS54*, increased in bvFTD (BA4-microglia) and *SLCO1A2,* a GWAS hit in PSP (Chen et al., 2018) increased in PSP (INS-microglia). Thus, disorder-specific DGE identifies genes previously associated with disease-relevant neurodegenerative phenotypes in bvFTD and PSP, as well as previous disease-associated microglial markers in AD (Wendimu and Hooks, 2022), which our cross-disorder design now shows were specific to AD.

### Patterns of shared and distinct composition of transcriptomic cell states across brain regions in dementia

We next systematically characterized the shared and distinct disorder-associated changes of 181 cell states across brain regions and different dementias to characterize new molecular markers and drivers of neuronal vulnerability, neuroinflammation and resilience. We used multivariate analysis (Methods) to rigorously identify changes in abundance of cell states across disorders within each brain region and cell type (Figure 2A, 2B, 2C).

**Figure 2:**
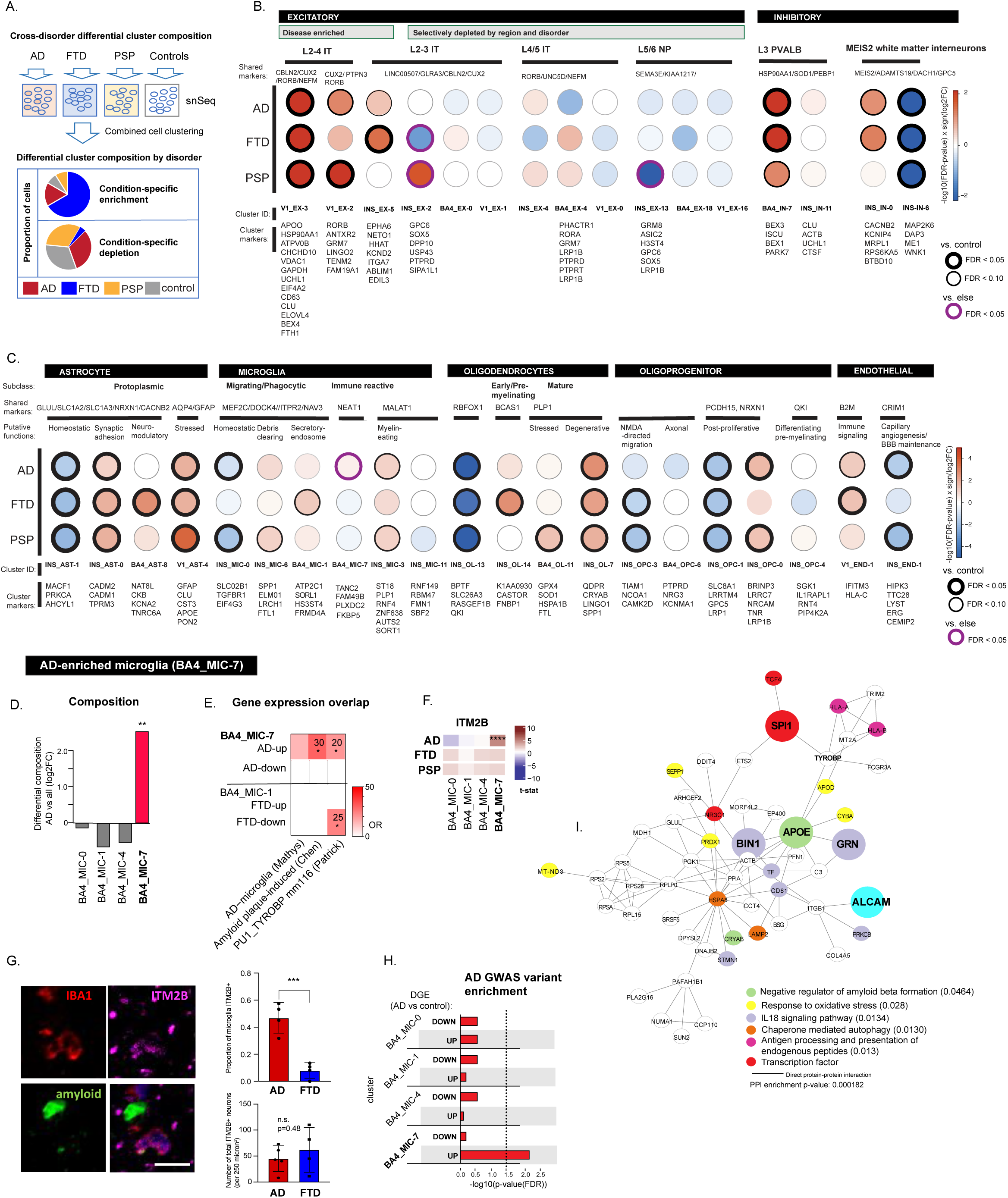
Shared and distinct neuronal and glial disease states in disorders versus controls. (**A**) Schema of analysis strategy for identifying shared and distinct disease-associate neuronal and glial states based on proportional differences in cluster composition of disease and control nuclei. (**B,C**) Heatmap of selected (**B**) neuronal and (**C**) glial clusters depicting the relative proportion of disease compared to control cells for each cluster and disorder shown, with colors ranging from red where disease cells are enriched, and blue where disease cells are depleted, based on compositional analysis (shown is log10(FDR) x sign(log2FC); with multiple testing correction applied within cell type and brain region, limma, Table S3), with boundary thickness and color indicating clusters with statistically significant enrichment (FDR<0.05 (thick) or FDR <0.1 (thin), comparing disease to control with multiple testing correction applied within cell type and region (black), or comparing one disease with all other conditions with multiple testing correction applied across brain regions (EX) or cell type and region (MIC) (see Table S3). Above each cluster are marker genes shared by related clusters based on hierarchical clustering (Figure 1, S2) and below are cluster-specific marker genes based on differential expression (Table S2), and putative associated functions based on gene ontology and literature review (Methods). <u>AD-specific microglia state from motor cortex (BA4_MIC-7),</u> showing (**D**) differential composition of BA4 microglia clusters in AD vs other samples (bvFTD, PSP, control) (Log2FC, limma, **FDR = 0.009, 4 comparisons), **(E**) Fisher’s exact test of overlap between genes differentially expressed (up or down regulated) in BA4_MIC-7 in AD cases, and BA4_MIC-1 in bvFTD cases (Table S5) compared to genes up-regulated in AD-associated microglia based on published reports (MIC1(Mathys *et al*., 2019), amyloid plaque associated microglia (Chen *et al*., 2020) and *PU.1* and *TYROBP* associated microglia module mm116 (Patrick *et al*., 2021) (Table S4), * FDR < 0.05 corrected for 12 comparisons. (**F**) Heatmap showing cluster-specific differential up-regulation compared to controls of *ITM2B* in BA4_MIC-7 microglia from AD cases, compared to all other BA4 microglia clusters and disorders (LME; t-stat shown; **** FDR<0.001, FDR corrected over 3,135 genes; Table S2B). (**G**) IBA1+ microglia demonstrate increased ITM2B protein staining in microglia in AD brains compared to bvFTD brains (frontal cortex, unpaired T-test, ***p = 0.0009, n = 4), but there is no change in the overall density of ITM2B stained neurons. (**H**) Enrichment of AD GWAS variants among genes up-regulated (up) in AD cases compared to control in BA4_MIC-7 microglia but not in genes down-regulated (down) or in other BA4 microglia clusters (MAGMA p-value with FDR over 8 comparisons). (**I**) Direct PPI network among genes up-regulated in BA4_MIC-7 in AD samples vs all other conditions (Table S5) highlighting in large circles AD disease genes and in colored circles genes that participate in significant gene ontology categories (FDR p-values for enrichment shown to right) as shown.

#### Shared disease-associated cell states in dementias

We observed 49 subclusters out of 181 (27%) that showed differential composition in one or more diseases (FDR <0.1), with nineteen depleted and thirty-one enriched in one or more disease conditions, the majority of which were observed across multiple disorders (66%; n = 33; Table S3). These shared disease-associated states include interneurons, astrocytes, microglia and OPCs that show robust changes in composition across all three disorders, and that have not been previously implicated in either AD, bvFTD or PSP (Figure 2B, 2C; Table S2, S3). A summary of clusters significantly enriched or depleted across disorders and brain regions and their markers is presented in Figure 2 and Supplementary Table 3, demonstrating a wide spectrum of glial and neuronal subtypes that are similarly depleted or enriched in brain samples from all disorders.

### Novel shared cross disorder changes in neuronal cell states

Taking advantage of robust markers from the BICCN reference atlas, we were also able to identify previously unknown robust disease-associated changes across multiple cell types and disorders (Summarized in Tables S3, S5). One such cell was a recently described population of white matter interneurons (Bakken et al., 2021) marked by *MEIS2*/*ADAMTS19* (Figure S3E, Figure 2B, Table S2, Table S3). These interneurons were spilt into two clusters in the insula, one composed predominantly of control cells (INS_IN-6), and one enriched for patient cells (INS_IN-0) (Figure 3E). The disease-enriched *MEIS2* neurons (INS-IN-0) showed transcriptional evidence of cell stress and injury response, including downregulation of DNA repair genes (ATM and ELOVL4) (Xiao et al., 2019) and upregulation of genes involved in protein folding and amyloid (Figure S3E). This reflects a previously unrecognized change in state between disease and controls, rather than an actual depletion of *MEIS2* interneurons, since the total number of *MEIS2* cells was not significantly different between cases and controls (Figure S3E).

**Figure 3:**
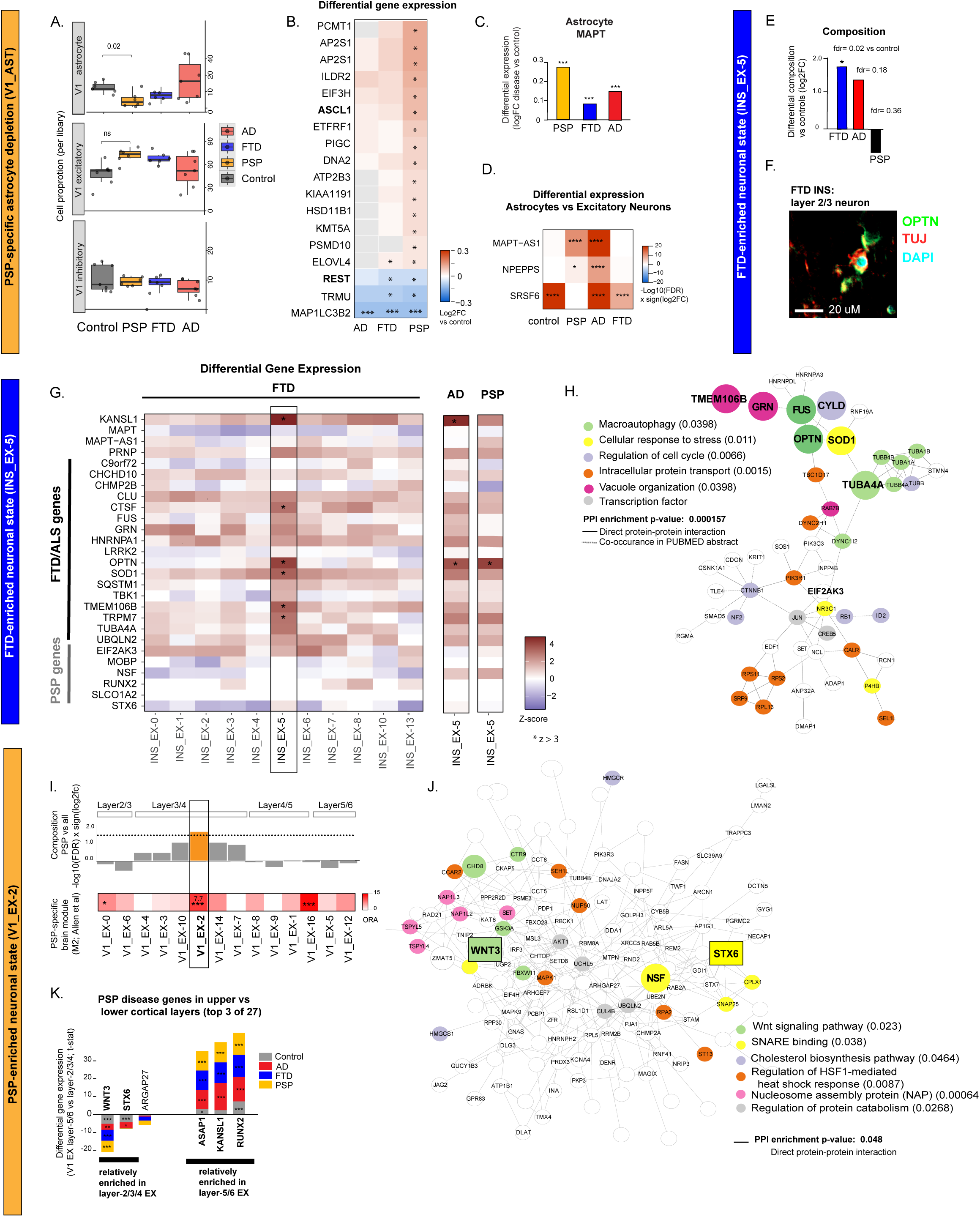
Disease-specific neuronal and glial states. <u>PSP-specific astrocyte changes</u> (**A**) Boxplot of cell proportions by subject in V1 for astrocytes, interneurons and excitatory neurons, showing significant depletion of astrocytes in PSP samples (limma with sample bootstrapping, FDR over 72 comparisons (3 diagnosis groups, 3 brain regions, 9 cell classes; Table S3). (**B**) Heatmap showing genes differentially expressed in PSP V1-AST (FDR<0.05 across 12677 genes, cross-disorder LME, Table S5). (**C**) Differential expression of *MAPT* in astrocytes (all regions combined) from AD, bvFTD and PSP compared to control cases (Table S5, Wilcoxin test, Seurat package, ***FDR<0.001 over 29 genes x 3 diagnosis groups = 87 comparisons). (**D**) Differential expression of *MAPT, MAPT-AS1*, *NPEPPS* and *SRSF6* in astrocyte compared to excitatory neurons within each disorder (all regions combined, Wilcoxin test comparison of mean percent cells with gene detected (Seurat, FindMarkers), n = 27; Table S5, quantified within subjects to eliminate possible effects of variable genetic background at the chromosome 17q21.31 locus). <u>bvFTD-enriched neuronal state from insular cortex (INS_EX-5)</u>, showing (**E**) differential composition vs control of nuclei in INS_EX-5 from bvFTD, PSP and AD cases (Log2FC, limma, *FDR = 0.02 over 11 comparisons Table S3). (**F**) OPTN protein staining of layer 2 excitatory neurons (TUJ1+) in bvFTD insular cortex. (**G**) Cross-disorder differential gene expression of bvFTD/ALS risk genes and PSP risk genes showing higher expression of multiple genes in INS_EX-5 neurons compared to other INS_EX clusters in bvFTD vs control cases; with AD or PSP vs control cases at right for comparison (Z-score, LME; * for Z -score > 3). (**H)** Functional PPI network among genes up-regulated in INS_EX-5 in bvFTD samples vs all other conditions (Table S5) highlighting ALS/bvFTD disease genes in large circles, and genes that participate in gene ontology categories with significant enrichment with colored circles as shown (FDR p-values for enrichment at right of each GO term). <u>PSP-enriched neuronal state in visual cortex (V1_EX-2),</u> showing (**I**) (top track) V1_EX-2 is enriched in nuclei from PSP cases vs other conditions based on differential composition analysis of V1-EX clusters (-log10(FDR p-value) x sign(log2FC) of PSP vs. all, p-value based on limma, Table S3, FDR over 14 comparisons), and (below track) comparison of PSP-brain specific modules described in Allen et al (Allen *et al*., 2018) with gene up-regulated in PSP vs control cases in each V1_EX cluster, showing significant overlap for clusters V1_EX-2, -16, and -0 (Fisher’s exact test, FDR correction over 14 comparisons) (* < 0.05, ** < 0.01, *** <0.001). (**K**) Stacked barplot showing differential expression (T-statistic, LME), measured within diagnosis group, of PSP risk related genes (Chen *et al*., 2018; Cooper *et al*., 2022) in layer 2/3/4 EX compared to layer 5 EX from calcarine cortex (LME, FDR * < 0.05, ** < 0.01, *** <0.001, n = 27,000 genes; Table S5). **(J)** Functional protein-protein interaction network that combines genes up-regulated in V1_EX-2 neurons in PSP samples (top 200 genes ranked by T-statistic >2, PSP vs all, LME, Table S5) with PSP disease genes enriched in layer 2/3/4 neurons (*WNT3*, *STX6*; in rectangle), indicating genes involved in gene ontology categories by color as shown (with FDR p-value for enrichment to right of GO term), and additional PSP disease genes by large size (Methods).

Another salient example involves *APOO*, previously reported as upregulated across multiple regions in AD (Liu et al., 2021). We identified a specific population of layer III/IV RORB+ excitatory neurons marked by *APOO* (V1_EX-3) that is significantly upregulated in the visual cortex in AD, bvFTD and PSP (Figure S3D). This distinct transcriptomic signature of *APOO* neurons from disease brain specifically overlaps with that of cultured IPSC-derived neurons expressing *MAPT* mutations compared to controls (Bowles et al., 2021) (Figure S3D). Since IPSC neurons are relatively immature (Gordon et al., 2021) and do not manifest neurodegenerative phenotypes *in vitro*, we hypothesize that these *APOO* neurons in vivo represent an early, relatively resilient neuronal disease state that is shared across disorders, given the relative sparing of neurons in visual cortex across all disorders (Braak et al., 2006; Kim et al., 2020; Kovacs et al., 2020).

### Novel shared cross disorder changes in glial cell states

We also observed shared changes involving glial states not previously associated with AD or any other dementia (Figure S2, Figure 2C). Notable examples include an astrocyte state depleted across disorders and regions and marked by *MACF1*, *PRKCA* and *AHCYL1* (INS_AST-1, BA4-AST-1, V1-AST-1) (Figure 2C, Table S2, Table S3) and cross disorder depletion of OPC populations (BA4_OPC-1; Figure 2C, Table S3) that are specifically marked by genes involved in injury response (Table S2). A population of microglia from the insular cortex was enriched in all disorders (INS_MIC-3) (Figure 2C, Figure S3F), overlapping significantly with disease-associated microglia described in MS brain (Figures S3G), suggesting that it may be a generalizable activated state. Other microglial states observed in insular cortex across disorders include depleted homeostatic states (Figure 2C, Figure S3F-G) and up-regulated states representing specific pathway perturbations (Figure S3H-I), including PI3K loss-of-function (LOF) (INS_MIC-0), Apoprotein and WNT LOF (INS_MIC-1), V-type ATPase LOF, PPAR-receptor agonist with WNT and PI3K pathway dependence (INS_MIC-3), or V type ATPase LOF with WNT LOF (INS_MIC-11) (Figure S3H).

We also reproduce previously observed changes in glial types in AD, including increases in *QDPR* OL (INS_OL-7, BA4_OL-6, V1_OL-4) and decreases in *PDE1A* OL (INS_OL-2), both of which are also shared with bvFTD and PSP (Figure 2C, Figure S2B, Table S3, Table S4). Among astrocytes, we observe reproducible downregulation of *SLC1A3* (Leng et al., 2021; Mathys et al., 2019), a marker of protoplasmic astrocytes, shared in all disorders and brain regions as a common feature of the total astrocyte pool (INS_AST, V1_AST, BA4_AST) (Figure S3C). Finally, we observe reproducible enrichment of *SEMA3E* OPC (INS_OPC-0) and depletion of *GPC5* OPC (INS_OPC-1) that are shared in all disorders and occur in proliferating OPCs in disease regions, including the insular cortex (Figure 2C, Table S3, Table S4) (Mathys et al., 2019).

### Disorder-specific changes in cell composition in dementias

Next, we analyzed clusters differentially enriched or depleted in a single disorder and region to leverage the differential vulnerability of each region in the different disorders. Using a strict statistical cut off, we identified three AD, eight PSP and six bvFTD clusters with disease-specific trends (Figure 2B, 2B, Table S3). We summarize these findings in Figure 2, including differential abundance and markers of shared and distinct neuronal and glial clusters. We then used these clusters to further define how cell reactivity and function among discrete classes and subclasses of cells can vary across disorders and regions with similar and different pathological burdens. Notable examples of disorder-specific cell states are described below.

#### AD-specific compositional changes in neuronal and glia cell states

We first assessed the extent to which our findings from AD reproduce previously reported cell states in AD snRNAseq studies (Grubman *et al*., 2019; Mathys *et al*., 2019; Olah *et al*., 2020; Otero-Garcia *et al*., 2022; Zhou *et al*., 2020). Reassuringly, we identify previously reported patterns of cellular responses to AD, showing that our data set is consistent with others in AD (Leng *et al*., 2021; Mathys *et al*., 2019). This includes changes involving excitatory neurons, interneurons, astrocytes, OL, OPC, and microglia in both AD datasets (Figure S3A-C, Table S4, Methods). We validate robust AD associated cell states and clarify their variable expression across brain regions with differing degrees of AD pathology and across AD, bvFTD, and PSP (Figure S3A-S3D, Table S3). This includes AD-associated depletion of layer 4/5 excitatory neurons marked by *RORB* and *NEFM* (BA4_EX-4) (Leng *et al*., 2021), which we now show are specific to AD and not observed in the other two disorders (Figure S3A). Among interneurons, we replicate the observed enrichment of a cluster of parvalbumin interneurons (INS-IN-10) in AD samples (Mathys *et al*., 2019) compared with controls that is greater than in bvFTD and PSP samples (Figure S3B, Table S3). This is of interest because disease affects involving interneurons have potential to alter neuronal circuit functions and affect connectivity, which has been reported in AD cases in the insular cortex (Zhou *et al*., 2010).

#### An AD-specific amyloid-associated microglia marked by ITM2B

We also observed changes in the motor cortex that have not previously been reported from snRNAseq of human AD brain tissue in regions with more advanced pathology. Of note, we observe a microglia cluster (BA4_MIC-7; Figure 2D, Table S3) whose signature genes share significant overlap with those of amyloid plaque-associated microglia in AD brain (Figure 2E) (Chen *et al*., 2020), and are further distinguished by their up-regulation of *ITM2B* (Figure 2F), a gene harboring mutations that cause a dominantly-inherited AD-like dementia (Ghiso et al., 2000). This cluster also corresponds to a highly AD risk gene-associated and age-associated microglial type from the ROSMAP database (Patrick *et al*., 2021; Figure 2E), confirming its AD association. We used IHC to validate that indeed *ITM2B* is more abundant in AD than bvFTD in microglia (Figure 2G), confirming its relative AD specificity. To further test the enrichment of genetic risk for AD within BA4_MIC-7, we performed LD-score regression of AD GWAS summary statistics, which indeed demonstrated that the upregulated genes in this AD-associated microglia were significantly enriched for common genetic risk variants (Figure 3H, Methods), including genes involved in amyloid processing and cellular response to oxidative stress (Figure 3I). The enrichment of genetic risk for AD within these BA4_MIC-7 genes supports their potential as therapeutic candidates.

### PSP- and bvFTD-specific compositional changes in neuronal and glia cell states

#### Astrocyte depletion in PSP

Canonical cell proportions were largely similar across disease conditions and brain regions at the level of major cell classes, and relative brain cell proportions also matched those reported in published reference data (Bakken *et al*., 2021). A singular, notable exception was the total count of astrocytes in the PSP brain. We observed that total astrocyte counts were significantly depleted in the visual cortex (FDR<0.05; astrocytes, Methods) of PSP cases compared with controls, which has not previously been recognized (Figure 3A). We therefore used two methods to robustly classify cells to confirm reproducibility including cell annotation by reference-based mapping (log2FC -2.04, FDR 0.07) and by published markers with bootstrap analysis (Methods; log2FC -1.29, FDR 0.023; Table S3). To consider the potential mode of PSP-specific astrocyte depletion, we performed disorder-specific differential gene expression and found 55 genes differentially expressed in astrocytes in the visual cortex from PSP cases compared with controls (FDR <0.1) (Figure 3B; Table S5, Methods). Of note was *REST*, which was downregulated specifically in PSP and bvFTD astrocytes in V1 (Figure 3B, Table S5). *REST* is a major regulator of the integrity of astrocyte-specific gene expression that suppresses neuronal gene expression in non-neuronal cells (Masserdotti et al., 2015). Similarly upregulated in PSP astrocytes was *ASCL1* (Figure 3B, Table S5), a chromatin remodeling factor that is sufficient to drive non-neuronal cells, including cultured astrocytes, towards a neuronal fate (Masserdotti *et al*., 2015) (Figure 3B). In both cases, the direction of changes in both of these major transcriptional drivers of cell fate is expected to reduce astrocyte-specific identity, which may contribute to their depletion in the PSP visual cortex, which has not previously been recognized.

### Increased expression of MAPT in PSP astrocytes

Astrocytic tau aggregation is a prominent feature of PSP (Chung *et al*., 2021), but it is not known how astrocytes differ in PSP compared to other disorders, including whether they are more vulnerable to cell death in PSP. We observed higher expression of *MAPT* mRNA in PSP astrocytes compared to controls, more so than for bvFTD and AD (Figure 3C, Table S5). In addition, astrocytes in PSP were further distinguished from those in AD and bvFTD by having lower expression of genes that reduce the expression of *MAPT* or its PSP-associated 4R isoforms. These genes reduced in astrocytes include *MAPT-AS1*, *NPEPPS,* a major cytosolic peptidase that directly degrades tau (Karsten et al., 2006;(Sengupta et al., 2006), and the *MAPT* exon 10 splice modifier, *SRSF6* (Yu et al., 2004) (Figure 3D, Table S5). These data indicate that accumulation of tau in PSP astrocytes likely involves post-transcriptional mechanisms.

### Excitatory neuron clusters specifically enriched in PSP and bvFTD

We also identified two clusters of excitatory neurons with significant disorder-specific enrichments, one in bvFTD (INS_EX-5; Figure 3E-H) and the other in PSP (V1_EX-2; Figure 3I-J). In bvFTD, in addition to the depleted cell class INS_EX-2 described below in the section on selective vulnerability, we observe enrichment in EX-5, which comprises a layer 2/3 excitatory neuronal cluster in the insular cortex (INS_EX-5) that was more significantly enriched in bvFTD compared to other disorders (Figure 3E, Table S3). INS_EX-5 neurons in bvFTD uniquely manifested increased expression of multiple bvFTD/ALS risk genes (LME, T-statistic >3), including *OPTN, TUBB1A, SOD1,* and *TMEM106B* (Figure 3G), and genes that function in associated pathways including macro-autophagy, stress response and vacuole organization (Figure 3H), which is especially remarkable because this is the most vulnerable cortical layer in the region in our study with the highest pathological burden (Kersaitis *et al*., 2004).

The PSP-enriched neuronal cluster was marked by *RORB/IL1RAPL1/UNC5D* in layer 3/4 excitatory neurons in the calcarine cortex (V1_EX-2) (Figure 3I, Table S3), a relatively spared region. This cluster overlaps with PSP-associated changes previously identified in bulk tissue from PSP (Allen et al., 2018) (Figure 3I), which we now localize to a distinct PSP-enriched neuronal cluster within superficial layer excitatory neurons. Pathway analysis of genes up-regulated identified cholesterol biosynthesis, *WNT* signaling, and synaptic vesicle cycle, including *NSF*, a PSP risk gene within the major 17q21.3 PSP risk haplotype (Chen et al., 2018; Sanchez-Contreras et al., 2018) (Figure 3J, Table S3, Methods); other PSP associated genes, *WNT3* and *STX6 (Chen et al., 2018; Cooper et al., 2022; Pittman et al., 2004)* were also significantly enriched in superficial neurons, compared to layer 5-6 neurons in control cells (Table S5; Figure 3K). These neurons in a relatively spared region, the visual cortex, appear to be up-regulating risk genes that may be related to their resilience (Figure 3J).

### Disorder-enriched glial states

We also observed several distinct glial states involving astrocytes, oligodendrocytes, and microglia, that are specifically enriched in one disorder, including AD-enriched microglia described above (BA4_MIC-7), bvFTD-enriched oligodendrocytes (INS_OL-14) and astrocytes (BA4_AST-8), and PSP-enriched oligodendrocytes (BA4_OL-11) (Figure 2C, Tables S2, S3; Figure S4A). In addition to the AD-specific microglia, BA4,MIC-7, we observed two other distinct disease-associated microglia clusters in BA4 (Table S3, Figure S4B-D). One population, BA4_MIC-4, was most abundant in the cases with higher tau pathology (Figure S4E). This population likely represents dystrophic microglia, showing high *FLT* and up-regulation of senescence-associated genes (Figure S4B,F). We also observed a microglia population most enriched in bvFTD (BA4_MIC-1; Figure S4G-J). This cluster was marked by *ATP2C*1, *SORL1*, *PLXDC2*, *HDAC9*, and tau modifier genes *HS3ST4* (Ferreira et al., 2022; Wang et al., 2020) and *FRMD4A* (Figure S2G, Figure S4B, Table S5) (Morita et al., 2017; Lambert et al., 2013; Yan et al., 2016). These *ATP2C1* microglia uniquely up-regulated pro-inflammatory signals (Guler et al., 2015; Freeman et al., 2017) (Figure S4H) and abundant in BA4 and INS, vulnerable regions in bvFTD (Table S3).

### Molecular markers of selectively vulnerable projection neurons in AD, bvFTD and PSP

We next mobilized the cross-disorder design to understand factors driving differential vulnerability of specific neuronal subtypes. We observed notable layer-specific depletion of disorder-specific neuronal subclusters in regions with moderate to high neuropathology scores (AD: BA4_EX-4; bvFTD: INS_EX-2; PSP: INS_EX-13; Figure 4A, 4B; Figure S5A; Table S3). The depleted populations we identified matched known expected patterns of vulnerability based on prior neuropathological data (Kersaitis *et al*., 2004; Leng *et al*., 2021) but have not been identified in PSP and have not previously been characterized at the molecular level in any disorder. As expected in bvFTD, the selectively depleted layer 2/3 cortical neurons (INS_EX-2; Figure 4A-B, Figure S5A) were found in the insular cortex and marked by layer-specific marker genes *CBLN2*, *CUX2*, and *RASGFR2* (Figure S5B) (Kersaitis *et al*., 2004). Similarly, in AD, the selectively depleted layer 4/5 EX neuronal population (INS_EX-4; Figure 4A-B, Figure S5A) was found in the motor cortex and marked by layer-specific marker genes *TSHZ2*, *FOXP*2, and *IL1RAPL2* (Figure S5B), together with previously reported markers of AD vulnerability, including *RORB* and *NEFM* (Figure S5B) (Leng *et al*., 2021; Zhou *et al*., 2020).

**Figure 4:**
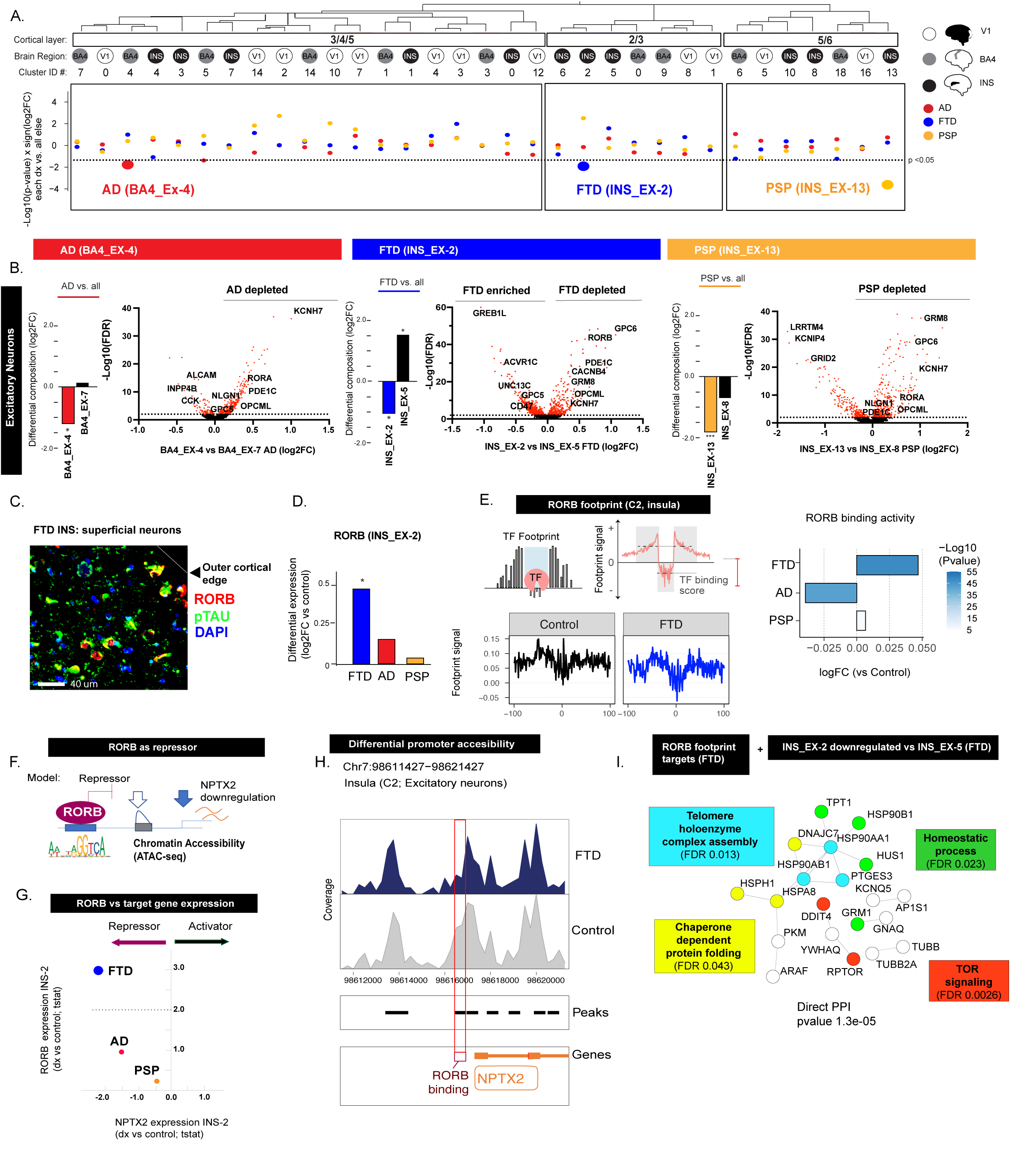
Cross-disorder comparisons of selectively depleted neuronal clusters identify *RORB* as shared repressor of disease-associated genes. (**A**) Differential cluster composition by diagnosis group across all excitatory neuronal clusters, based on proportion of nuclei from each diagnosis category relative to all other groups combined. Clusters group by cortical layer, which is indicated, below which brain region of origin is labeled and colored (black =insula; grey = BA4; and white = V1). Below each cluster, colored by disorder (AD in red, bvFTD in blue, PSP in yellow), is the differential composition score (Methods; -log10(FDR) x sign (log2FC)) for each cell type cluster, showing one different depleted clusters for each disease. (**B**) Differential composition in diagnosis indicated vs all other samples (left, limma, *FDR<0.1 over 2 comparisons) and differential gene expression within diagnosis group (right, LME; Methods) comparing each disorder-specific selectively depleted cluster with a matched cluster that is not depleted in disease, but shares overlapping subclass-specific markers (see Figure S4C; Table S5; FDR correction > 23,365 genes). (**C**) RORB immunostaining in layer 2/3 cortical neurons in bvFTD insular cortex. (**D**) Differential expression of *RORB* in disease vs control cells from INS_EX-2 (LME, *FDR<0.05 corrected over 3 disorders). (**E**) Differential *RORB* binding in layer 2/3 excitatory neurons (ATAC INS_EX subcluster C8) in bvFTD, AD and PSP vs controls based on chromatin footprinting (Methods). (**F**) Model showing *RORB* as a candidate repressor of *NPTX2* in selectively depleted neurons. (**G**) Differential expression of *RORB* relative to *NPTX2* in INS_EX-2 neurons in bvFTD vs control, AD vs control and PSP vs control samples (LME, Table S5). (**H**) Chromatin accessibility peaks based on snATAC-seq at the *RORB* binding site in the *NPTX2* promoter of INS_EX (ATAC cluster C2). (**I**) Protein-protein interaction plots highlighting genes with enriched gene ontology among genes relatively downregulated in bvFTD-depleted neurons relative to other layer 2/3 neurons (INS_EX-2 vs INS_EX-5, bvFTD samples, t<-2, LME, Table S5) that are also bvFTD-specific *RORB* target genes based on chromatin footprinting (bound in bvFTD but not control, PSP or AD samples) (showing direct protein-protein interaction (PPI) p-value, and gene ontology enrichment p-values from String (Methods)).

In PSP, where cortical laminar vulnerability patterns have not been fully established (Ohm et al., 2022), we observed a selectively depleted population of a layer 5/6 near-projecting neuronal cluster (INS_EX-13; Figure 4A-B, Figure S5A-B). Based on brain connectivity maps (Gehrlach et al., 2020; Ghaziri et al., 2018), these near-projecting (NP) neurons in the insula may innervate nearby subcortical structures with high tau pathology burden in PSP (Kovacs *et al*., 2020), including the striatum and globus pallidus. We next assessed whether these layer 5/6 NP neurons differentially express genes involved in PSP pathogenesis compared to other layer 5/6 excitatory neurons. Indeed, we identify expression of PSP risk genes, including *RUNX2, STX6*, *MOPB* and *EIFAK3* (Chen *et al*., 2018; Sanchez-Contreras *et al*., 2018) in this class of vulnerable neurons (Methods, Figure S5C, Table S5).

To further investigate the enrichment of PSP risk variants in this vulnerable class of INS_EX-13 neurons, we analyzed 27 genes whose expression was recently shown to be regulated by functionally validated PSP genetic risk variants that underlie the major PSP GWAS signals (Chen *et al*., 2018; Cooper *et al*., 2022). We compared expression of the risk genes implicated by these variants between layer 2/3/4 and layer 5/6 neurons, finding evidence of significant DGE of multiple PSP risk genes, with the majority enriched in deep layer neurons compared to superficial layer neurons, including *RUNX2*, *KANSL1*, *ARL17B*, *MAPT*, *ASAP1*, *LINC02210-CRHR1* and *SP1* (13 out of 15 risk genes significantly different at FDR<0.05, test, Table S5). This indicates that known PSP risk genes are enriched in layer 5/6 near projection neurons that are depleted specifically in PSP cases relative to controls (INS_EX-13), supporting a causal link between PSP genetic risk (Table S5) and this uniquely vulnerable cell population identified by cross-disorder differential cell composition analysis (Figure 4A-B, Figure S5A-B, Table S3).

Disorder-specific depleted neurons expressed additional genes other than their canonical marker genes that distinguished them from non-depleted neurons in the same cortical layer (Figure 4B). We first compared markers of selectively depleted neurons across disorders and found examples of both disorder-shared and disorder-distinct genes (Figure 4B, Figure S5D). *KCNH7, OPCML, PDE1C* and *NLGN1* were notable as shared markers of selectively vulnerable neurons in bvFTD (INS_EX-2, Layer2/3 IT), AD (INS_EX-4, Layer4/5 IT) and PSP (INS_EX-13, Layer5/6 NP). Remarkably, a recent study identifies *OPCML* and *KCNH7* (Kv11.3) as genes conferring resilience in mouse lines and nominates *KCNH7* as a druggable target (Telpoukhovskaia et al., 2022) and *PDE1C* and *NLGN1* are known drivers of neuronal toxicity (Hollerhage et al., 2017; Kattimani and Veerappa, 2018). We extend these observations to human brain here and further validate that *KCNH7* marks vulnerable, depleted neurons identified in an independent dataset in AD (Leng et al., 2021). *GRM8* (Sanchez-Juan et al., 2014) and *GPC6* are shared markers of PSP- and bvFTD-depleted neurons, *RORA* of PSP- and AD-depleted neurons, and *RORB* of AD and bvFTD depleted neurons (Figure 4B-D). However, AD vulnerable neurons express high levels of *RORB* at baseline in controls (Figure S3A), whereas bvFTD-depleted neurons only up-regulate *RORB* in disease cases and not in controls (Figure 4D), suggesting that *RORB* may be an inducible marker of bvFTD-vulnerable neurons. Lastly, in AD-depleted neurons, *GPC5* was up-regulated, which we validated using IHC, showing that *GPC5* is a new marker of AD vulnerable neurons that has not been previously recognized (Leng et al., 2021; Mathys et al., 2019). *GPC5* shows increased expression in AD relative to controls, localizes to the surface of layer 4/5 neurons where it is depleted in AD cases, and it colocalizes with hyperphosphorylated tau (Figure S5E).

### snATAC seq validates RORB as a shared transcriptional driver of selective vulnerability in AD and bvFTD

Our analysis shows that *RORB*, a previously established marker of neuronal vulnerability in AD (Leng *et al*., 2021), was also upregulated in bvFTD-depleted neurons (Figure S3A, Figure 4C-D). To experimentally confirm if *RORB* TF binding activity was increased in excitatory neurons from the insular cortex in bvFTD samples, we performed snATACseq in insular cortex of 9 bvFTD, 8 control, 10 AD and 11 PSP subject samples, and conducted chromatin footprinting of *RORB* targets among selectively vulnerable layer 2/3 excitatory neurons in bvFTD (Methods; Figure 4E). As expected, *RORB* gene regulatory activity, as measured by gene promoter occupancy, was higher in bvFTD cases compared to controls in INS_EX-2, Layer 2/3 IT neurons, but not in AD or PSP samples (Figure 4E). Thus, *RORB* expression and its activity is increased in bvFTD vulnerable INS_EX-2, Layer 2/3 IT neurons.

We further explored the functional implications of increased *RORB* activity in selectively depleted neurons by broadening the analysis to all annotated *RORB*-target genes that show differential RORB binding in open chromatin peaks of selectively vulnerable excitatory neurons in bvFTD and AD. We found two genes, *TMEM196* and *NPTX2*, whose promoters include a *RORB* binding motif and showed reduced chromatin accessibility and down-regulated gene expression in vulnerable neuronal populations in both disorders (bvFTD samples, layer 2/3 depleted neurons (INS_EX-2 vs EX-5); AD samples, layer 4/5 depleted neurons (BA4_EX-4 vs EX-7); Figure 4H, Table S5, Table S7). Consistent with our prediction that RORB represses *NPTX2* expression (Figure 4F) we observe an inverse correlation between *RORB* expression and *NPTX2* expression across disorders within this class of layer 2/3 neurons (Figure 4G; INS_EX-2 and INS_EX-5; Methods). This finding is notable because *NPTX2* functions in synapse homeostasis (Galasko et al., 2019) and is a prognostic biomarker whose expression anti-correlates with disease progression in AD and bvFTD (Libiger et al., 2021; van der Ende et al., 2020). In bvFTD, we find that the majority of the annotated *RORB* targets were primarily downregulated in selectively vulnerable neurons (Methods; 120 downregulated, 14 upregulated, FDR < 0.05; INS_EX-2 relative to INS_EX-5 in bvFTD). These down-regulated genes form a highly significant PPI network (direct PPI enrichment p-value 1.3E-05; Methods) reflecting a coordinated stress response, and its down-regulation here indicates a relative dampening of this protective pathway (Figure 4I). In contrast, upregulated *RORB*-bound target genes in bvFTD-depleted neurons included *BICC1*, an RNA binding protein that controls protein synthesis and stress granule formation (Table S5) (Estrada Mallarino et al., 2020; Iaconis et al., 2017)). These findings support a potential for *RORB* to promote neuronal vulnerability in bvFTD by driving deleterious alterations in the coordinated cellular stress response and RNA solubility.

### Transcriptomic drivers of shared and distinct disease-associated cell states

The identification of *RORB* as a potential driver of neuronal vulnerability in both AD and bvFTD suggested that analysis of additional TF mediated drivers of disease-associated cellular states would be of value. Moreover, identification of gene regulatory networks (GRNs) places DGE as components of known coherent biological processes (Network, 2021). We used single-cell regulatory network inference and clustering analysis (SCENIC) (Figure 5A, Methods; Aibar et al., 2017; Polioudakis et al., 2019) to identify cell-type specific GRN regulons for excitatory neurons, astrocytes, oligodendrocytes, and microglia, and define the disease specificity of cell-type specific regulons (Figure 5A-B, Figures S6A-C, Table S6) We validated snRNAseq based SCENIC predictions of cell type- and disease-specific regulon activity by snATAC-seq, assessing TF binding site enrichment as well as TF footprinting in 78 disease and control samples (Methods), focusing our analysis on excitatory neurons (114,6872 nuclei) and microglia (15,715 nuclei) (Figure 5D-E, Figure S6 D-E, Table S7, Table S1).

**Figure 5:**
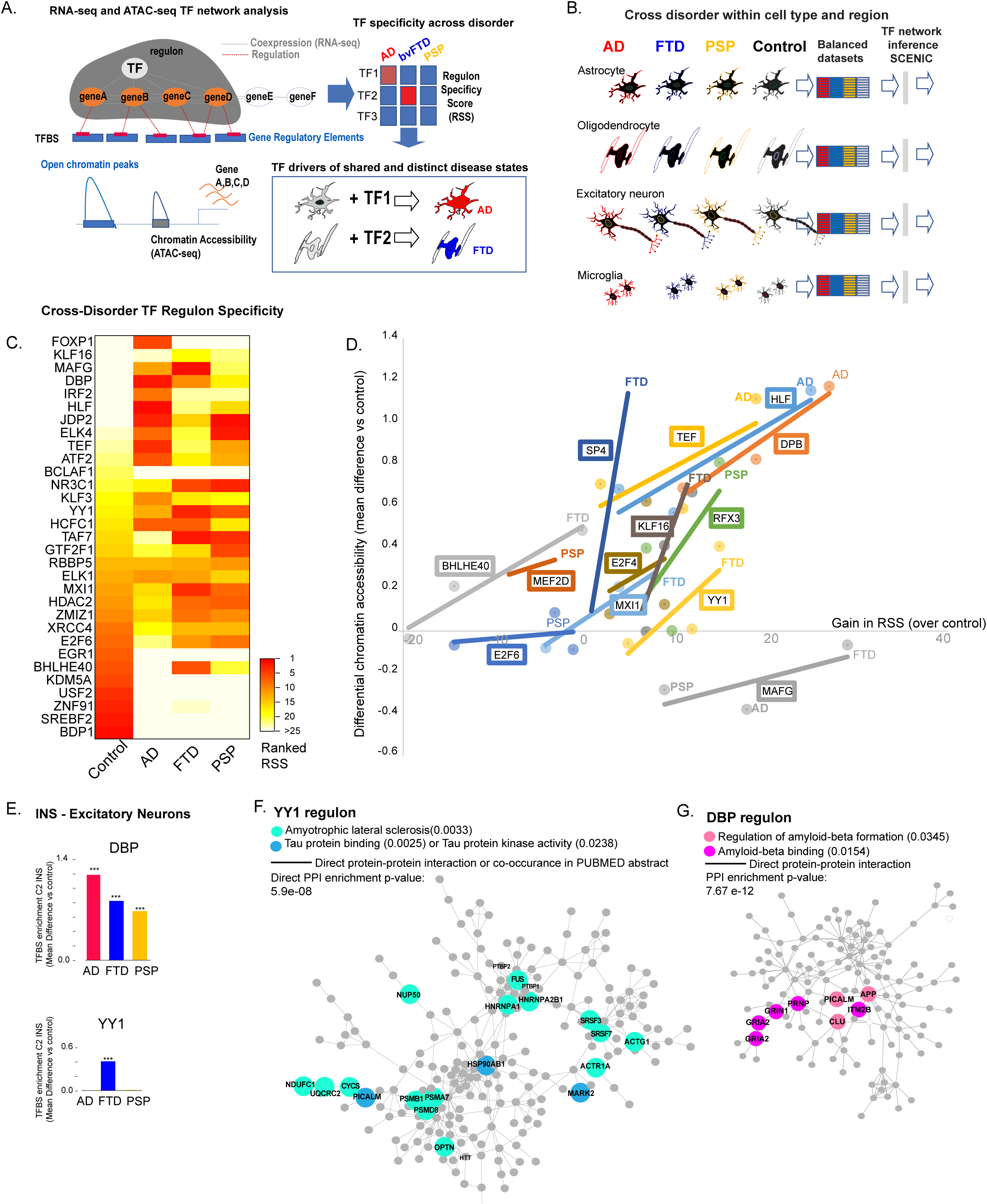
Transcription factor network inference (SCENIC) identifies transcription factor regulons active across cell types, disorders and brain regions. (**A**) Workflow for validation of SCENIC TF results using snATAC and chromatin footprinting. (**B**) Schema of strategy for cross-disorder comparison of TF activity within cell type and brain region. (**C**) Heatmap comparing INS EX neuron regulon specific scores across disorders, showing relative rankings across disorder of the 10 regulons per disorder. (**D**) Scatterplot of validated disease trends among top 10 most disease associated TF for each disorder, based on regulon specificity score, by comparing cross-disorder differences in regulon specificity score (gain in rank vs control) vs cross disorder differences in differential chromatin accessibility (refers to Table S6). (**E**) Validation of differential chromatin accessibility at *DBP* and *YY1* binding sites in AD, bvFTD and PSP vs control INS EX (C2; ChromVar) compared to ranked regulon specificity score (RSS rank (SCENIC) for AD, bvFTD, PSP; respectively; *DBP* ranked 2, 10, 18; *YY1* ranked 13, 3, 6). (**F,G**) Direct PPI and associated pathways enriched among (**F**) *YY1* and (**G**) *DBP* regulon target genes from INS EX (STRING).

First, we empirically defined 1332 cell context-specific regulons driven by 250 TFs, 65% of which were active in only one cell type, including TFs well known to be cell type-specific, such as *TBR1* and *SCRT1* in EX (16 total unique); *SOX9* and *NFAT* in AST (82 unique) and *PRMD1*, *SPI1*, *TAL1* and *IRF8* in MIC (57 unique) (Table S6, Figure S6A-C). More than half of the 10 most active TF regulons in each disorder were highly active in multiple disorders (Table S6). We highlight the 10 most specifically active TF regulons in each disease (Figure S6C), which revealed a core set of TF regulons that were consistently active in multiple disorders compared to controls in more than one cell type or brain region, including TFs previously implicated in neurodegenerative diseases such as *CEBPB* (Tittelmeier et al., 2020) (Dent et al., 2021), *YY1*, a regulator of cell death (Chen and Chan, 2019), *HDAC2*, an AD associated gene whose suppression reverses cognitive decline in AD models (Gediya et al., 2021), and *NFE2L1*, a neuroprotective gene (Katsuoka et al., 2022). In contrast, TF regulons more active in control samples have known functions in supporting learning and memory and cellular maintenance (*CREB*, *EGR1*, *SOX8*) (Turnescu et al., 2018); Figure S6C, Table S6).

In addition, we identified TF regulons more specific to one disorder and or region (Figure S6C, Table S6), particularly among astrocytes and microglia where over 50% of the top 10 active TFs were exclusive to one diagnosis group. In microglia in particular, distinct clusters of TFs co-varied by diagnosis and brain region (Figure 6A), suggesting distinct TF networks likely contribute to the diverse microglial transcriptomic states observed in this dataset (Figure S2G).

**Figure 6:**
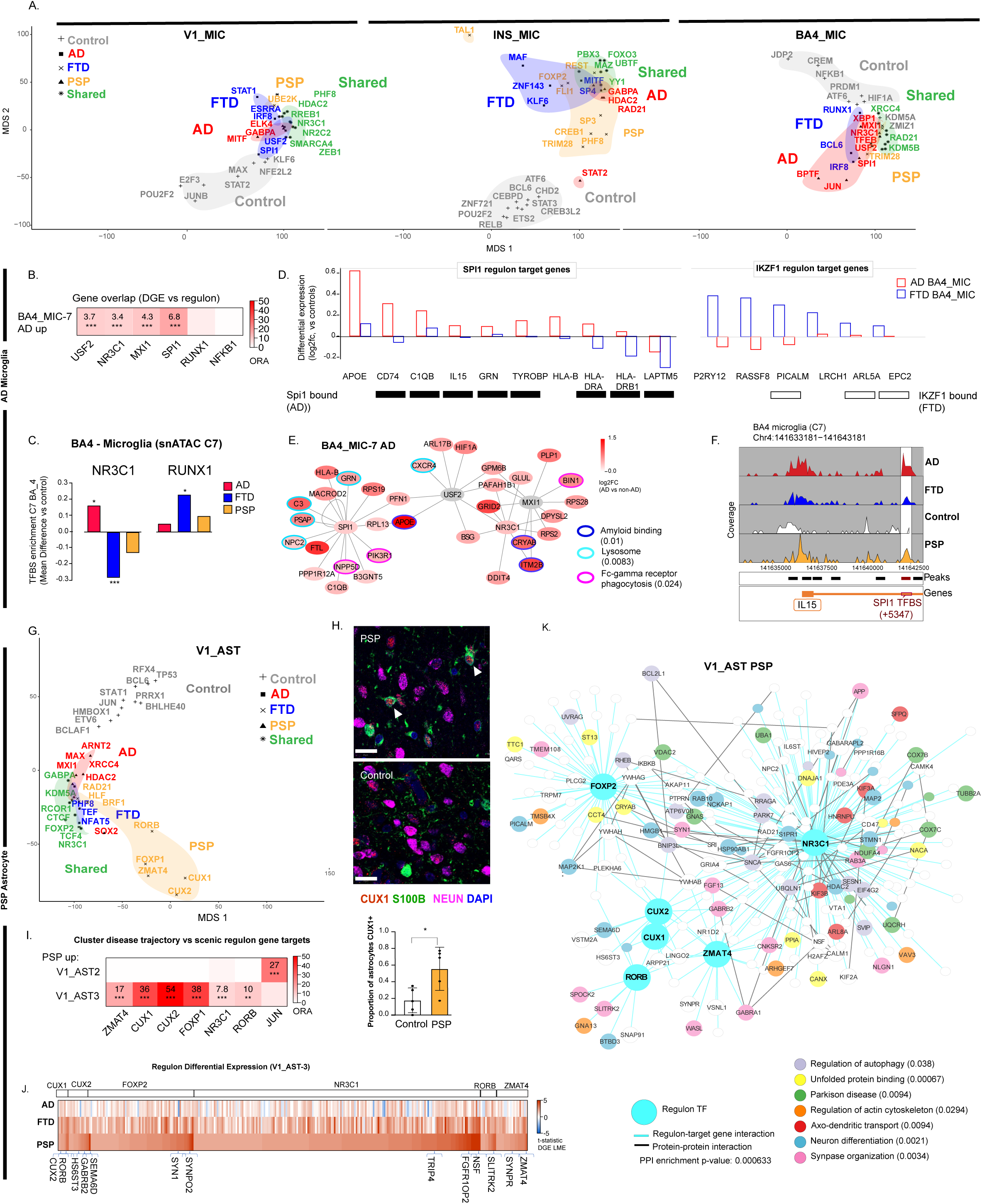
Distinct transcription factor networks drive disorder-specific glial states in AD and PSP. (**A**) Multidimensional scaling plots using Sammon projections (Methods) showing relatedness of transcription factor regulons specific score ranks across diagnosis within microglia (MIC) comparing three brain regions, and highlighting distinct and shared TFs across disorder based on ranked differential entropy score (Methods). (**B**) Gene overlap enrichment in AD BA4 MIC between specific regulons and genes up-regulated in AD-associated BA4_MIC-7 (T-statistic >2, LME), comparing regulons with high regulon specificity for AD microglia (*USF2* (ranked 2) *NR3C1* (ranked 4), *MXI1* (ranked 6), *SPI1* (ranked 7) and low regulon specificity for AD microglia (*RUNX1* (ranked 44) and *NFKB1* (ranked 48)) (Fisher’s exact test, ***FDR<0.001 over 5 comparisons). (**C**) Validation of differential chromatin accessibility at *NR3C1* and *RUNX1* binding sites in AD, bvFTD and PSP vs control BA4 MIC (C7; ChromVar, *FDR<0.05 over 453 TFs measured per disorder) compared to ranked regulon specificity score (SCENIC; for AD, bvFTD, PSP; respectively: *NR3C1* ranked 4, 9, 7, *RUNX1* ranked 13, 3, 6,). **(D)** Differential expression of *SPI1* and *RUNX1* regulons target genes BA4 MIC comparing AD and bvFTD samples vs, controls (LME; Table S5). Black boxes below each *SPI1* target gene indicated binding in AD samples based on chromatin footprinting (BA4 C7; Table S7). White boxes below each *RUNX1* target gene indicated binding in bvFTD samples based on chromatin footprinting (INS C7, Table S7). (**E**) TF regulon plot showing target genes for *USF2*, *NR3C1*, *MX11* and *SPI1* regulons upregulated in AD BA4_MIC-7 with edge length proportion to (1-GRN) score (Table S6) and node color proportion to differential expression in AD BA4_MIC-7 (LME, AD vs all conditions, Table S5, Methods). Key indicates specific genes in major enriched GO pathways with FDR corrected p-values shown to their right. (**F**) Differential chromatin accessibility (peak density) at *SPI1* binding site on *IL15* promoter in AD compared to bvFTD, PSP, control BA4 microglia (BA4 C7, snATAC, Methods). (**G**) Multidimensional scaling plots using Sammon projections (Methods) showing transcription factor regulons specific score ranking across diagnosis within AST from V1 (Table S6). (**H**) Immunohistochemistry showing *CUX1* in PSP and control astrocyte and neurons, with quantification showing great percent of astrocytes positive for *CUX1*+ in PSP V1 samples (unpaired T-test, *p = 0.023, n = 5). (**I**) Heatmap showing that TF regulons with high and low specificity for PSP V1 astrocytes (relative specificity score RSS) significantly overlap with genes specifically up-regulated in PSP-in V1_AST-3 astrocytes (T-statistic > 2, LME) (Fisher’s exact test, FDR applied over 14 comparisons, **<0.01,***<0.001). (**J**) Heatmap showing higher expression in PSP V1_AST-3 astrocytes of regulon target genes of *ZMAT4*, *RORB*, *NR3C1*, *FOXP2*, *CUX1* and *CUX2* (T-statistic, LME, Table S5). (**K**) Combined PPI and TF regulon network plots, showing targets of *ZMAT4*, *RORB*, *NR3C1*, *FOXP2*, *CUX1* and *CUX2* regulons from V1-AST with direct PPI and their significantly enriched GO terms, including PD and unfolded protein binding.

We confirmed increased chromatin accessibility at TF binding sites with high regulon specificity scores (Figure 5D-E, Figures S6D-E; Methods). Across each of the top 10 TFs for INS_EX, we observed a consistent correlation across disorders between ranked differences in regulon specificity and chromatin accessibility (Figure 5D-E). Notable ALS/bvFTD disease genes *OPTN* and *FUS* were targets gene of *YY1* (Figure 5F), whose regulon activity ranked highest in in bvFTD (Figure 5C-E), while AD disease genes *APP*, *ITM2B* and *PICALM* were target genes of *DBP* (Figure 5G) whose regulon ranked highest in AD samples (Figure 5C-E).

Next, we identified combinations of regulons that reflected the majority of the gene expression difference associated with disorder-specific clusters (Figure S7A-D, Figure 6B), including *RELA* and *NELFE* in bvFTD INS OL (INS_OL-14), *ATF4* and *SOX10* in PSP BA4 OL (BA4_OL-11), *SOX5 and NFAT5* in bvFTD astrocytes (BA4_AST-8), *RUNX1* and *IKZF1* in bvFTD microglia (BA4_MIC-1), and *SPI1* and *NR3C1* in AD microglia (BA4_MIC-7) (Figure 6B-D). We confirmed disorder-specific increase in chromatin accessibility at binding sites of *RUNX1* in bvFTD and *NR3C1* in AD (Figure 6C-D). Orthogonal experimental data from gene knockout studies allowed us to independently confirm that both *RUNX1* and *IKZF1* are functional drivers of the bvFTD-enriched *ATP2C1* microglia signature (BA4_MIC-1; Table S6, Methods; (Subramanian et al., 2017)). Furthermore, validated *IKZF1* regulon target genes were upregulated in bvFTD microglia relative to controls (Figure 6D). These data support a potential role for *RUNX1* and *IKZF1* in bvFTD-specific reactive microglial states observed in a brain region with moderate bvFTD pathology (BA4_MIC-1).

### AD disease genes engage combinatorial TF programs regulating AD-specific microglia

Of interest was an AD-specific microglial TF network comprised of *SPI1*, *NR3C1*, *MXI1* and *USF2* based on high regulon specificity, and high overlap with gene expression changes; their combined targets were responsible for a remarkable 60% of up-regulated genes in the amyloid associated, AD-specific cluster (Figure 6B). Therefore, we examined the AD-specific TF program identified in microglia to understand the combinatorial regulation of disease-specific microglia states both at the RNA and chromatin-level. For example, we observed that *SPI1*, which is known to govern the expression of AD disease genes in myeloid cells (Pimenova et al., 2021), drives expression of a distinct gene set involved in lysosome and Fc-gamma receptor mediated phagocytosis, including multiple AD risk genes including *APOE, INPP5D, GRN,* and the AD biomarker, *IL15* (Figure 6D-E, Figure S7E) (Popp et al., 2017; Wightman et al., 2021). We validated the AD specific increase in *IL15* promoter accessibility at the *SPI1* binding site by footprinting (Figure 6F). In contrast, the genes regulated by another TF driver, *NR3C1,* participate in amyloid processing and microglial reactivity (Figure 6E; (Baik et al., 2019)), including the marker gene *ITM2B*, the marker of BA4_MIC-7 in AD brain (Figure 2G; Table S7), which we validated by footprinting (Table S7). This demonstrates that the effects of *NR3C1* and *SPI1* direct discrete, biological pathways that act combinatorially to modulate the BA4-7 microglial neuroinflammatory state in AD.

### Astrocytes in PSP ectopically express neuronal TFs and disease related pathways

As described above, a prominent pathological feature of PSP is astrocytic tau inclusions (Chung *et al*., 2021). We observed that astrocytes in PSP downregulate *REST* (Figure 3B) and found that four predominantly neuronal TFs appeared to be uniquely active in astrocytes in PSP: *CUX1*, *CUX2*, *ZMAT4*, and *FOXP1* (V1_AST-3, Figure 6G). As *CUX1* regulated genes are typically neuronally expressed (Cubelos et al., 2015; Rodriguez-Tornos et al., 2016), we tested the prediction that it was ectopically expressed in astrocytes in PSP. We confirmed that CUX1 selectively stains neurons in control brain tissue, but robustly stains S100b-positive astrocytes in PSP samples (Figure 6H). In PSP, the V1_AST-3 cluster gene expression signature significantly overlapped with these regulons (Figure 6I) and showed disease-specific up-regulation of *ZMAT4* (log2FC 0.24, FDR 0.023, Table S5) and a broad trend of increased expression of their targets (Figure 6J). These targets include several Parkinson’s Disease (PD) disease genes, such as *PARK7* and *SNCA* (Corti et al., 2011), regulators of autophagy and the unfolded protein response (Figure 6K). Therefore, ectopic expression of neuronal transcription factors specific to PSP astrocytes underlie PSP-specific transcriptomic changes in changes involving both neuronal genes and known causal pathways in PD.

### MAFG-NFEL2L1 regulates PSP disease genes and neuroprotective pathways in excitatory neurons that are selectively dampened in PSP vulnerable neurons

It was intriguing that our GRN analysis identified *NFE2L1* and *MAFG* as manifesting higher activity in more vulnerable neurons, because *NFE2L1* and *MAFG* are known to cooperate to regulate gene programs involved in proteostasis, antioxidant response and xenobiotic stress factors (Katsuoka *et al*., 2022). Evidence for *NFE2L1* activity was markedly increased in all disorders in visual cortex, which is relatively spared from neurodegeneration in all disorders (Figure 7A). *NFE2L1* and *MAFG* regulons both included genes involved in iron sequestration (*FTH1* and *FTL*), and the protective ALS/bvFTD gene *SQSTM1* (Figure 7B, C, Table S6). In addition, the MAFG regulon includes *VCP*, another ALS/bvFTD disease gene that reduces tau proteopathic seeding (Zhu et al., 2022), the PD risk gene *SCNA*, and genes involved in the regulation of selective autophagy, proteostasis, RNA stabilization, and synaptic vesicle exocytosis (Figure 7D). We confirmed that *VCP* expression is positively correlated with *MAFG* expression across EX in the insular cortex (Figure 7E). We validated several targets by direct chromatin footprinting, including *UBE2N*, *KIF5C*, *RARB* and *VCP* (Figure 7D, Table S7). These findings suggest a model whereby the *MAFG/NFE2L1* regulon protects neurons from tau aggregation (Figure 7F). Indeed, we find that tissue samples containing more neurons with high regulon activity have lower tau pathology scores (cor -0.57 p=0.04; Methods, Figure 7G). Moreover, neurons that are selectively depleted in one disease also lose their regulon activity in a disorder-specific fashion (Figure 7H), consistent with the known function of *MAFG*/*NFE2L1* in cellular resilience.

**Figure 7:**
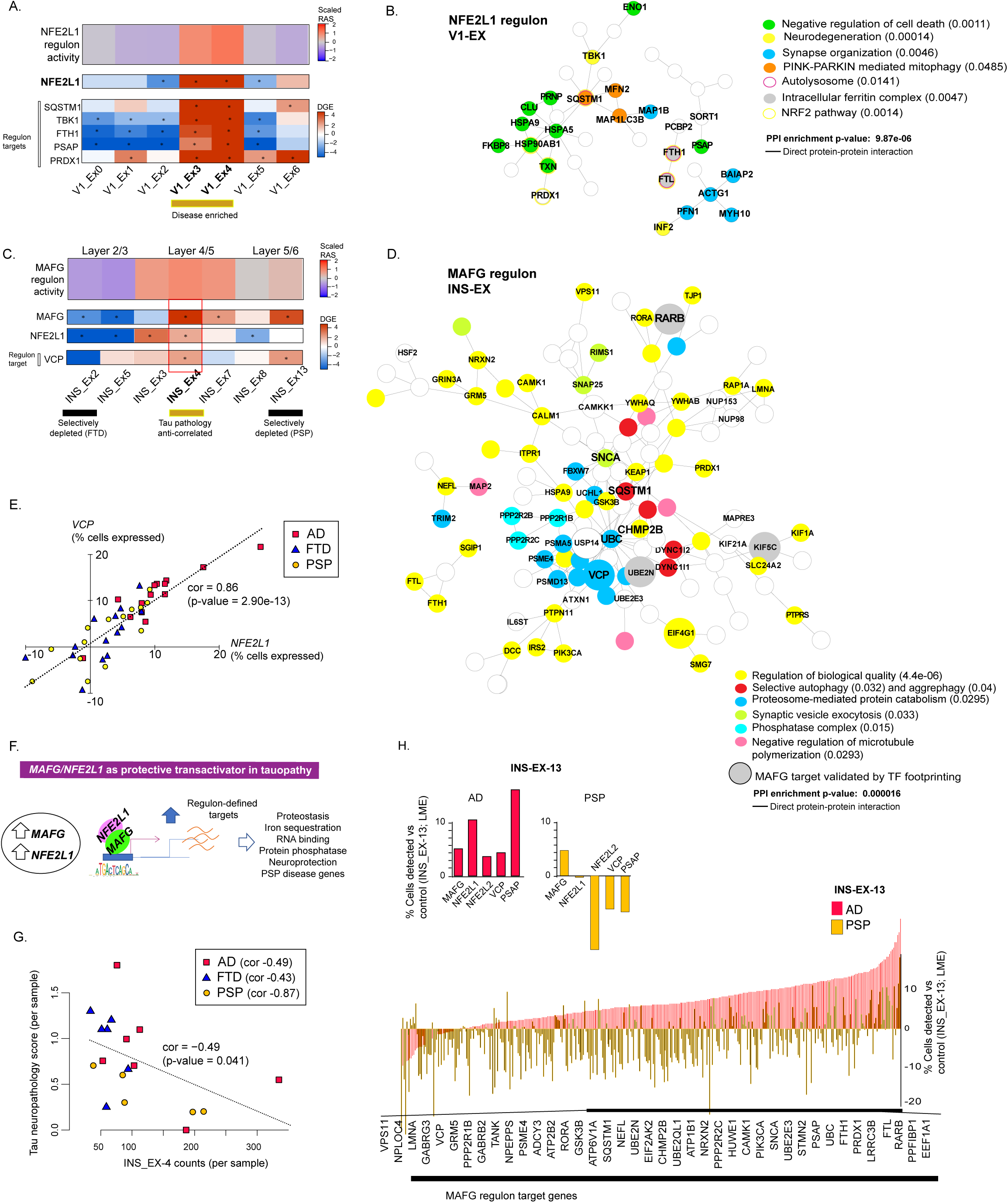
*MAFG/NFE2L1* drives a resilience program of proteostasis across disorders with inversed expression relative to selective neuronal loss. (**A**) Cross-cluster comparison between *NFE2L1* regulon activity score (colored by scaled RAS, Key; see Methods, SCENIC) and the expression of *NFE2L1* and selected target genes in V1 EX neurons (each cluster *vs* all remaining V1-EX neurons; LME, Methods; * T-statistic > 2; Key = T statistic, Table S2). (**B**) Direct PPI plot of *NFE2L1* regulon (top 250 genes) showing genes participating in enriched disease related gene ontology pathways, such as mitophagy and cell death (STRING, FDR of enrichment versus genome as background). (**C**) Cross-cluster comparison among INS EX of *MAFG* regulon activity score (see Methods SCENIC)), and the expression of *MAFG* and selected target genes (each cluster vs all remaining INS-EX neurons; LME, * T-statistic > 2; Key = T statistic, Table S2). (**D**) Direct PPI plot of the *MAFG* regulon (top 250 genes) showing genes participating in enriched gene ontology pathways (STRING, FDR of enrichment versus genome as background). (**E**) Scatterplot showing change in expression of *NFE2L1* compared to *VCP* in each INS_EX cluster across PSP, AD, and bvFTD samples relative to controls with trendline (Pearson’s correlation, p-value = 2.9e-13, n = 42). (**F**) Model showing a neuroprotective gene program induced by *MAFG/NFE2L1* complex in cells otherwise vulnerable to neurodegeneration, including PSP disease genes. (**G**) Scatterplot and correlation trendline showing sample level counts of INS_EX-4 neurons compared to Tau neuropathology score (Pearson’s correlation and p-value calculated across all disorders shown with trendline. Correlations calculated within each diagnosis are also shown in key. (**H**) Differences in percent cells expressing *MAFG*, *NFE2L1*, (left) select target genes, and (right) the entire *MAFG* regulon in INS_EX-13 neurons comparing AD and PSP samples.

## Discussion

Several studies have illustrated the power of single cell RNA sequencing to elucidate pathways dysregulated in AD (Grubman *et al*., 2019; Mathys *et al*., 2019; Morabito *et al*., 2021; Otero-Garcia *et al*., 2022; Sadick *et al*., 2022; Zhou *et al*., 2020), but whether these changes are specific to AD is not known. By employing a cross disorder design and by profiling multiple regions with differential vulnerability, we characterize both cross-disorder and disease specific pathways underlying differential susceptibility and resilience. Our analyses strongly support the value of direct cross disorder comparative analysis for defining disorder-associated cellular trajectories. Notably, we were able to replicate several previous observations in AD single cell data including the association of *RORB* with AD vulnerable neurons and disease effects involving interneurons, OPCs, and astrocytes, providing additional confidence that our data and analyses are robust. We also identify multiple novel disease-specific homeostatic and pro-inflammatory states, including depleted astrocyte populations marked by *MACF1*, shared disease effects involving changes in the state of *MEIS2* white-matter interneurons, shared depletion of *ENPP6/RBFOX1* expressing oligodendrocytes (Hughes and Stockton, 2021; Xiao et al., 2016), and OPC populations marked by genes involved in migration and NMDA sensing (Xiao *et al*., 2013). We annotate shared and distinct disease states comprehensively to provide an extensive resource, which we use to understand the molecular features and transcriptional drivers of selective vulnerability and resilience, including the role of disease-specific risk genes. We identify specific transcriptional programs representing coherent biological processes that define the single cell pathophysiology of these disorders. By coupling snRNAseq with ATAC-seq we experimentally validate the bio-informatically predicted regulatory drivers of these altered cell states.

Despite recent progress (Leng *et al*., 2021; Roussarie et al., 2020), the molecular basis of selective neuronal vulnerability in neurodegeneration is relatively uncharacterized. Such factors might be divergent in different disorders, or shared. Here, we identify four genes that are reproducibly identified in multiple classes of depleted neurons across all three disorders, including *OPCML* and *KCNH7*, which have also been identified experimentally in mouse models of AD (Libiger et al., 2021; van der Ende et al., 2020). Given the consistency of this observation, we interpret these to be shared components of differential neuronal vulnerability. Other components of neuronal resilience across disorders include upregulation of the transcriptional regulators, *MAFG/NFE2L,* which coordinate the proteostatic stress response (Katsuoka *et al*., 2022) including the expression of *VCP*, an bvFTD/ALS risk gene (Ling et al., 2013). Although known to be neuroprotective, the relationship of *MAFG/NFE2L* to vulnerability in neurodegenerative dementias was not previously known. The upregulation of *MAFG/NFE2L1* and their targets with binding to cis-regulatory regions confirmed by single cell TF footprinting, in specific spared populations of excitatory neurons in AD (INS_EX-3,4,7,13), bvFTD (INS_EX-3,7,13) and PSP (INS-EX-3,4,7), illustrates how cell intrinsic differences in injury response may also be a major determinant of vulnerability in dementia. This includes differentially affected neuronal subtypes that match expected patterns of selective vulnerability, such as those harboring gene regulatory programs driven by *RORB* in AD and bvFTD depleted neurons, and *GRM8*, a candidate genetic risk modifier for CJD (Sanchez-Juan *et al*., 2014), in PSP- and bvFTD-depleted neurons. Moreover, we were able to identify the specific molecular programs underlying superficial projection neuron vulnerability in bvFTD and identify for the first time an additional class of projection neurons, INS_EX-13, selectively depleted in PSP. These pathways that vary across spared and dying/depleted cell populations become potential therapeutic targets based on the supposition that restoring or boosting resilience-associated factors present in spared neurons may be protective (Karsten *et al*., 2006).

In this regard, glial subtype heterogeneity in the brain is increasingly appreciated (Chen *et al*., 2020; Endo et al., 2022; Grubman *et al*., 2019; Mathys *et al*., 2019; Olah *et al*., 2020; Rexach *et al*., 2020; Sadick *et al*., 2022; Zhou *et al*., 2020). But how this heterogeneity corresponds to specific neurodegenerative conditions is not understood. We replicate previous findings of a dystrophic microglial state in post-mortem brain from patients with AD (Nguyen *et al*., 2020; Olah *et al*., 2020). We also find evidence of microglia heterogeneity relative to pathology and disorder, including an amyloid-associated microglia state unique to AD (BA4_MIC-1), a bvFTD-enriched state with up-regulation of genes involved in sterile inflammation (BA4_MIC-1;(Freeman et al., 2017) and a cross disorder state with high levels of WNT signaling genes (INS_MIC-3). This implicates both shared and distinct neural immune pathways in each disorder, which may specify disorder-specific therapeutic targets.

We also uncover previously unrecognized glial diversity correlated with the degree of tau pathology, such as BA4_MIC-1 and INS_MIC-1). Furthermore, we defined and validated a combinatorial TF gene regulatory network underlying the AD specific microglial state, BA4_MIC-7, that includes *SPI1*. It is notable that a recent study demonstrated that *SPI1* represses AD risk genes in late-stage AD (Morabito *et al*., 2021), and is a risk locus whose reduced expression is associated with delayed AD onset (Huang et al., 2017), nominating it and downstream pathways as potential therapeutic targets. Disease associated glial states involve differential expression of genes that have been associated with AD or pathological tau burden, including AD risk genes in AD-specific microglia (*ITM2B*, *APOE*) and two modifiers of tau burden (*FRMD4A*, *HS3ST4* (Ferreira et al., 2022; Wang et al., 2020; Yan et al., 2016)) marking microglia enriched in bvFTD brain. These data support suggest that disease-specific genetic risk variants influence microglial states observed in the brain in patients with these dementias, which in turn differentially modulate distinctive aspects of AD and bvFTD pathology.

In the case of PSP and bvFTD, we observe distinct changes involving astrocyte and oligodendrocyte clusters V1_AST-3 in PSP and INS_OL-14 in bvFTD. PSP is pathologically defined by its unique pattern of neuronal and astrocytic tau pathology (Chung *et al*., 202; (Roemer et al., 2022), but its molecular correlates have not been well-defined. In PSP, we find that astrocytes (V1_AST-3) are uniquely depleted in the visual cortex where they down-regulate *REST*, which represses neuronal gene expression, while upregulating transcription factors typically observed in neurons, including *CUX2* and *ZMAT4 (Cubelos et al., 2015; Weed et al., 2019)*, as well as *MAPT* itself. We hypothesize that the loss of factors driving astrocyte identity or self-maintenance is related to their reduction in the primary visual cortex in PSP; which is consistent with the trend towards increased neurons in the primary VC (Table S3). Since this is the first time this has been observed in PSP, this observation enabled by unbiased molecular profiling is exciting, but warrants additional confirmation studies.

Tying molecular phenotypes such as gene expression to causal factors requires understanding how they are related to genetic risk. In this regard, we provide several examples of AD, bvFTD, or PSP risk genes that are enriched in specific cell types that are preferentially observed in that disease, suggesting a causal link. This includes AD risk genes such as *ITM2B*, *APOE*, *BIN1*, and *SPI1* in microglia that are enriched in AD brain, bvFTD/ALS risk genes including *OPTN*, *CTSF, TPRM7* and *TMEM106B* enriched among layer2/3 neurons that are specifically vulnerable in bvFTD (INS_EX-2), and PSP associated genes such as *WNT3* and *RUNX2* that were differentially expressed in layer 2/3 neurons (e.g. V1_EX-2) that are more abundant in PSP, or layer 5/6 neurons that are depleted in PSP (e.g. INS_EX-13). These data show that known risk genes act in specific neuronal and glial states or cell types that differ across related disorders, primarily non-neuronal cells in AD and specific neurons in bvFTD and PSP. Moreover, causally associated disease states are not broadly distributed in the brain, but rather are limited to specific cell types and brain regions. This further underscores the importance of examining multiple brain regions to understand causal disease pathways at the cellular level, which we show can provide a clearer picture of cross-disorder and disease-specific aspects of resilience and vulnerability, thus informing the therapeutic roadmap.

## Acknowledgements

Funding for this work was provided by Roche Pharmaceuticals (D.H.G., D.M.), BrightFocus (D.H.G., J.E.R), Rainwater Charitable Foundation (D.H.G. and W.W.S), NIH grants (K08 NS105916 (J.E.R), R01 AG075802 (J.E.R., L.T.G), and John Douglas French Alzheimer’s Foundation (J.E.R.). The UCSF Neurodegenerative Disease Brain Bank is supported by NIH grants AG023501 and AG019724, the Rainwater Charitable Foundation, and the Bluefield Project to Cure bvFTD. The University of Pennsylvania Center for Neurodegenerative Disease Research is supported by NIH grant P01AG066597, P30AG072979 and U19AG062418

## Author Contributions

J.E.R. and D.H.G. designed and supervised all RNA, ATAC and IHC experiments, analyzed and interpreted results, and wrote this manuscript. D.M. contributed to the design of the RNA experiments, RNA sequencing, and revision of the manuscript. J.E.R. completed the experiments, performed the analyses and generated figures and tables with additional contributions of co-authors to experiments as follows. Y.C. performed single nuclear ATAC data analysis including chromatin footprinting and provided technical training and supervision of the ATAC analyses. L.C. provided technical assistance with analysis of single nuclear ATAC and RNA data including data processing, hierarchical clustering and differential gene expression, differential cluster composition, ATAC data filtering and clustering experiments and TF enrichment analyses from ATAC. D.P. performed single nuclear RNA data processing including sample filtering, clustering and subclustering with Liger. L-C.L. and W.W.S. advised regarding subject and region selection. L.C-L., W.W.S. and J.R. curated brain tissues for analysis. L-C.L. and S.E.G prepared tissue specimens. W.W.S., L.T.G., and S.S. performed regional neuropathological assessments and scoring and, with J.Q.T. and E.B.L, performed neuropathologic diagnostic assessments. V. M performed immunohistochemistry experiments. A.E. performed SCENIC analysis and doublet removal. A.Y assisted with nuclear isolation and 10x reactions. D. C. contributed to library sequencing. R. K. performed major cell type composition analysis with bootstrapping. J. O. generated RNA and ATAC libraries. J. H. assisted with nuclear isolation. H. V. curated and provided disease and control brain tissue slides for IHC in independent samples.

## Methods

### Human Tissue Samples

Freshly frozen human brain tissue (BA4: precentral gyrus, V1: calcarine cortex, INS: insular cortex) were obtained from the UCSF Neurodegenerative Disease Brain Bank and University of Pennsylvania Center for Neurodegenerative Disease Research Brain Bank. We obtained samples from 40 total individuals, including 10 subjects with clinical diagnosis of bvFTD and neuropathological diagnosis of Pick’s disease (FTLD-tau), 10 subjects with clinical diagnoses of AD-type dementia and a neuropathological diagnosis of Alzheimer’s disease, and 11 subjects with a clinical diagnosis of PSP-RS and a neuropathological diagnosis of PSP (FTLD-tau), and and 10 non-demented controls, sex-matched to the patients (Table S1). All procedures involved the use of postmortem human brain were conducted after obtaining the written informed consent, and approved by the Committee on Human Research at the University of California San Francisco and University of Pennsylvania. IRB exemption was obtained from the UCLA IRB to authorize use of de-identified human postmortem brain single nuclear sequencing data in this study. Neuropathological diagnoses were made prospectively at the contributing brain banks following standard criteria(Mackenzie et al., 2010; Montine et al., 2012). Of 120 samples used as input for snRNA-seq, 118 passed nuclear isolation of which 12 failed library synthesis and 5 were removed as sample outliers (Table S1) leaving 101 final samples. For snATAC-seq, 80 samples yielded 78 final libraries post quality control, filtering and outlier removal (Table S1).

### Neuropathological scoring and brain region selection

Patients autopsied at UCSF underwent a standardized, semi-quantitative scoring of various pathological features (Table S1). These assessments were carried out prospectively, at the time of autopsy. Frozen tissue blocks were taken from the same regions that underwent scoring; these regions were taken from an adjacent frozen slab whose tissue face fell across the plane of the dissection blade from the scored region. Scoring for neurodegenerative features and tau burden were carried out using methods previously described (Lin *et al*., 2019). Briefly, morphological and immunohistochemical analyses of glial and neuronal tau pathomorphologies were performed in 40 regions of 13 cases with progressive supranuclear palsy (PSP), 5 with Pick’s disease (bvFTD with tau pathology), and 20 with Alzheimer’s disease. Nonspecific features of neurodegeneration were scored based on the hematoxylin and eosin stain and included microvacuolation, astrogliosis, and neuronal loss, each graded on a 0 to 3 scale (absent, mild, moderate, severe). Tau aggregates were visualized using a monoclonal anti-phospho-tau (pS202) antibody CP13. Pathomorphologies were assessed using the same 0-3 scale and included neurofibrillary tangles, Pick’s bodies, neuronal cytoplasmic inclusions, globose tangles, astrocytic plaques, tuft-shaped astrocytes, thorn-shaped astrocytes, tau-positive threads and grains in the gray and white matter, and glial cytoplasmic inclusions. To analyze the pattern of tau-related pathomorphologies of the three patient groups, we calculated a composite score by adding tau and neurodegeneration scores. The composite score was used to prioritize brain regions from mild to severe stage of tau inclusions in each patient group of tauopathies. Overall, three cortical brain regions were selected across three patient groups to reflect selective regional vulnerability of tauopathies for this study, including middle insula (INS), precentral gyrus (BA4), and calcarine cortex (V1).

### Immunofluorescence

IHC-P human brain slides were deparaffinized by being placed in a Clarity™ oven for 15-30 minutes then submerged in Citrasolv (Cat# c9999, Sigma) to graded ethanol washes. Primary antibodies were tagged with Alexa Fluor® fluorescent secondaries, Alexa Fluor™ 555 Tyramide SuperBoost™ Kit (B40923) or Opal™ system fluorophores (NEL861001KT). All slides were treated with 0.3% Sudan Black in 70% EtOH for 2 min to reduce autofluorescence before imaging. Sections prepared for Alexa Fluor™ antibodies were heated in 1x Citrate Antigen retrieval (Cat #c9999, Millipore Sigma) before blocking (5% Donkey serum cat#NC1697010, Fisher) for 1 hour at room temperature. Primary antibodies (CUX1-1:150, Cat #ab54583, abcam; S100beta-1:2000 Cat #ab41548, abcam; NeuN-1:100, Cat #266004) were incubated at 4°C overnight. Secondary antibodies (Donkey anti-mouse 488, Cat#A-21202, Invitrogen; Donkey (Dk) anti-Rabbit 555, Cat#A31572; Goat anti-guinea pig 647, Cat#A21450) were diluted 1:500 and incubated for 1 hour. GPC5 staining used tyramide amplification, wherein sections were incubated in 3% hydrogen peroxide for 1 hour at room temperature and washed (1X PBS Cat#10010049, Fisher) before blocking (C2-kit) for 1 hour at room temperature. GPC5 (1:250 Cat# ab124886, abcam) primary incubation was followed by an HRP secondary incubation for one hour and a Tyramide amplification step. Sections were then stripped and reheated for 12 minutes in 1x Citrate Antigen retrieval. After rinsing with 1X PBS, a blocking step (5% Dk Serum) for 1 hour followed and then a 1 hour incubation of primary antibodies or counterstain (Phospho-Tau (AT8)-1:100, Cat#MN1020, Fisher; nissl N-21479-1:20). Secondary antibodies (Donkey anti-mouse 488, Cat#A-21202, Invitrogen.) were diluted 1:500 and incubated for 1 hour. For ITM2B staining and quantification we used the OPAL system, wherein slides were heated in kit’s (Antigen Retrieval pH6) buffer and directly placed in blocking buffer after 1X PBS rinses. Each primary antibody incubation (ITM2B-1:200 Cat# PA5-31441, Invitrogen; Iba1-1:200, Cat# ab5076, abcam; Bioss; IBA1 (ab5076), and amyloid (1:200 ab201060)) was preceded by 10 minutes of kit’s blocking buffer, followed by rinses in TBST, 10 minutes in secondary HRP and a ten minute RT incubation in respective opal fluorophore (480, 520, 570, 620, 690) diluted to 1:100 in appropriate 1x Amp diluent. DAPI (Anti-fade Cat#H-1800-10, VECTASHIELD®) was used to stain the nuclei.

### Image Quantification

All imaging and data analysis was completed in a blinded fashion. For CUX1 quantification, images were taken with upright scope using Zeiss software. The number of CUX-positive and S100beta-positiveF cells were blindly quantified in PSP and control cases (n=5) using a series 5 randomly selected, representative regions magnified at 20X. For ITM2B and GPC5 quantification analysis, we used the Vectra® Polaris™ microscope to scan slides with Opal™ system-tagged fluorophores antibodies and remaining stained sections. Using Qupath (v0.2.0-m5), 4 images from representative 40X magnified regions were quantified to measure ITM2B positive, IBA1 positive cells and total number of ITM2B positive DAPI positive cells, using across AD cases (n =4). To quantity GPC5 positive neurons, nissl staining was used to first distinguish upper vs deep cortical layers, as well as neurons. From a series of randomly selected images, we quantified the number of GPC5 positive neurons in layer ⅘ in AD and control cases (n=7,6), as well as the proportion of neurons staining for hyperphosphorylated tau (AT8).

### Single nucleus isolation

Nuclei were prepared from 60–70mg of frozen brain tissue per sample, with all procedures carried out on ice or at 4°C with RNase-free reagents. Briefly, postmortem frozen brain tissue was gentle lysed in 3mL homogenization buffer (250mM sucrose, 150mM KCl, 30mM MgCl_2_, 60mM Tris, 0.01% v/v Triton X-100, 0.001% v/v Digitonin, 0.01% v/v NP40, 1µM DTT, supplemented with 0.2U/mL RNase Inhibitor (NEB, M0314), Complete protease inhibitor cocktail (Roche, 11697498001)) using a Wheaton Dounce Tissue Grinder (30 strokes with pestle B). The lysate was filtered through a 40µm cell strainer and centrifuged at 1000xg for 8 minutes to obtain a nuclear pellet. To remove debris, the nuclear pellet was resuspended in 350µL homogenization buffer and 1:1 with an equal volume of 50% iodixanol buffer (Iodixanol 60% v/v combined with buffer of 250 mM sucrose, 150mM KCl, 3mM MgCl_2_, 60mM Tris), then layered over 600µL of 29% iodixanol buffer (Iodixanol 29% v/v combined with buffer of 250mM sucrose, 150mM KCl, 3mM MgCl_2_, 60mM Tris) and centrifuged at 13500xg for 20 minutes. The supernatant was discarded, and nuclei gently resuspended and washed in 1mL of 1% BSA/PBS. The nuclei were visually inspected to confirm complete lysis and nuclear integrity. Nuclei were manually counted and diluted to a concentration of 1000 nuclei/µL in 1% BSA/PBS. For single-nucleus RNA sequencing (snRNA-seq), libraries were prepared using the Chromium Single Cell 3’ Reagent Kits (v2 for BA4 and V1, v3 for INS) according to the manufacturer’s protocol (10X Genomics). For snATAC-seq, libraries were prepared using the Chromium Single Cell Next GEM Single Cell ATAC kit (v1.1). RNA-seq libraries were sequenced on a Novaseq S2 or S4 sequencer with paired end reads (read 1: 26 bp, read 2: 96 bp) targeting over 50,000 paired reads per nucleus. ATAC-seq libraries were sequenced on a Novaseq S4 with paired end reads (2x50 bp) targeting 25,000 paired reads per nucleus.

### Single nucleus RNA-seq Alignment and Filtering

Raw single-nuclei RNA-seq data was processed using the 10X Genomics Cell Ranger (v3.0) pipeline. Reads were aligned to the Ensembl release 93 *Homo sapiens* genome. Cells were selected for downstream analysis using the cell barcodes associated with the most UMIs. We estimated the number of cells expected to be captured based on input nuclei concentration and retained this many cell barcodes for downstream analysis. Cells with < 200 unique genes detected were removed (gene detection: > 1 count). Cells with > 8% of their counts mapping to MT genes were removed. Genes detected in < 3 cells were removed. Normalization was performed using Seurat (v3.1(Butler et al., 2018). Briefly, raw counts are read depth normalized by dividing by the total number of UMIs per cell, then multiplying by 10,000, adding a value of 1, and log transforming (ln (transcripts-per-10,000 + 1)). Raw UMI counts data were assessed for the effects from biological covariates (clinical diagnosis, anatomical region, donor, age, sex), and technical covariates (RIN, PMI, library batch, number of UMI, number of genes detected, percentage MT). The effects of the number of UMI (sequencing depth) were removed from the read depth normalized expression values using a linear model. Outlier samples were identified based on abnormal frequencies of major cell types and divergent gene expression patterns, and were removed from the analysis (Table S1). Doublet cells were removed using Doublet Finder (McGinnis et al., 2019). As doublets should be limited to within one sample library, each library was run independent scaling, principal component analysis, and clustering through Seurat. pK was estimated using paramSweep_v3 on the first 40 principal components. The homotypic doublet proportion was initially estimated at 7.6% and refined through modelHomotypic. Per-library doublet determination was performed using doubletFinder_v3.

### Annotation of major cell types, subtypes and states

All samples (BA4, INS, V1; all diagnoses) were jointly clustered in Seurat (v3.1). Each cluster was then annotated as a major cell type using mean expression of groups of cell type marker genes. Canonical genes were selected based on mouse and human studies as well as published reference atlas enriched genes (Astrocyte *GFAP*/*SLC1A2*/*AQP4*/*SLC14A1*, Endothelia *VWF*/*CLDN5*/*FLT1*, Ependymal *ARMC3*/*SPAG6*/*DNAH11*, Inhibitory neurons *GAD1*/*GAD2*, Excitatory neurons *SATB2*/*SLC17A7*/*NRGN*/*SNAP25*, Oligodendrocyte *MOBP*/*MOG*/*TF*, Microglia *CD47*/*CSF1R*/*C3*, OPC *VCAN*/*PDGFRA*/*CSPG4*, Pericyte *ACTA2*/*RGS5*/*PDGFRB*, Lymphocyte *CD247* taken from (Hodge *et al*., 2019; Kelley *et al*., 2018; Mathys *et al*., 2019; Sweeney *et al*., 2016) also confirmed all major cell type classifications used human motor cortex reference dataset using Azimuth (Bakken *et al*., 2021), a web-based portal from the Allen Brain Atlas (https://azimuth.hubmapconsortium.org). Neuronal layer-specific markers were assigned based on (Lake et al., 2018).

After identification and clustering of 9 main cell types, we sub-clustered each of the 9 main cell types independently to identify distinct cell states within each cell-class. To maximize the influence of disorder-specific effects on subclustering, we performed this analysis on each brain region separately, generating 134 total clusters (pre-filtering. Specifically, we combined nuclei from the same major cell type and different diagnosis groups, and performed batch correction, data integration and subsampling using Liger (v0.4.2; with kappa = 20 and lambda = 0.5) (Liu et al., 2020; Welch et al., 2019). We removed clusters that were non-representative across multiple subjects (with less than three libraries contributing >10 nuclei from at least one diagnostic group), low-quality clusters based on significant association with multiple sample quality metrics (FDR <0.05, limma, number of gene detected per cell, sample post-mortem interval, percent_mitochondrial genes detected per cell), or ambiguous cell type with 30% or more nuclei in a given cell-type enriched cluster bearing markers more suggestive of a disparate cell type (see Table S1 for full list of filtered clusters with justification).

For reference based assignment of clusters to cell classes and subclasses, cells were mapped to an external human motor cortex dataset (Bakken *et al*., 2021) using the reference-based mapping workflow described (Hao et al., 2021). We prefiltered our cells as recommended by the web interface: UMI count in the range [212, 33185], gene count in the range [201, 6486], and proportion of mitochondrial genes <= 2%. For visualization Sankey plots (Figure S1E) were made between our subclusters and the predicted subclusters. High gene count (nFeature_RNA >= 10000) and cells with low mapping scores (mapping.score <= 0.9) were filtered out. Subcluster/predicted subcluster links were restricted to those in the same cell type. Clusters that did not map to reference cell type were discarded from further analysis (Table S1). In this way, each final cluster was assigned a reference cell class and subclass, as shown (Table S2, Figure 1, Figure S1-S2). Finally, to organize subclusters into related cell types, subtypes and states across all brain regions and diagnoses groups, we performed hierarchically clustered based on their marker genes, and then jointly annotated based on marker gene expression and reference cell type assignments. First, for each cluster, we calculated cluster-specific marker gene expression by performing differential gene expression to compare each cluster with all other clusters of the same cell type and brain region. Genes were filtered using a minimum proportion, keeping only genes detected in 10% or more of cells per cell type x brain region group. Differential gene expression (DGE) was calculated based on mixed effects model used lmerTest::lmer (Kuznetsova, 2017) with formula expression ∼ clinical_dx + pmi + age + sex + number_umi + percent_mito + (1 | library_id). To complete hierarchical clustering across brain regions, we then filtered DGE lists to genes detected in each region. We then used the estimate terms (beta) for the top 100 genes per cluster based on DGE results for all regions, sorted by the cross-region variance. The top 100 most variable genes were used to compute a Euclidian distance matrix (stats::dist) and complete-linkage hierarchical clustering (stats::hclust) using default parameters. Groups of clusters were annotated based on shared marker genes. Reference-based cluster assignment results from Azimuth were overlaid manually to the hierarchical clustering framework to assign cluster groups to reference cell type and subtypes based on high confidence matches where >70% of cells of a cluster assigned to the same reference cell type (Table S2, Figure 1G, Figure S1-S2).

### Cell type composition analysis

Cluster composition was defined as the proportion of cells in a given cluster relative to the total number of cells of that major cell class and brain region, per subject. To measure cell type composition across subjects and diagnosis groups, cluster composition percentages were used as pseudobulk counts, forming a cluster by subject count matrix for each major cell type and brain region. The matrix was normalized using TMM (Robinson and Oshlack, 2010) and Limma-voom (Law et al., 2014), then fit on model formula ∼0 + dx + pmi + age + sex + mean_percent_mito + median_genes. T-statistics were calculated using eBayes (Smyth, 2005) and FDR-adjusted p-values were reported. For various analyses as reported, we then measured and reported different contrasts between samples, such as for one diagnosis versus control samples, all disease versus control samples, or one diagnosis versus all other samples. For selected clusters of interest, to correlate cluster counts with variation in neuropathology and tau scores across subjects (Table S1), we measured the Pearson’s correlation and p-value between cluster count proportion and neurodegeneration and tau score per subject across disease samples. Control samples were excluded because their tau and neuropathology scores were either zero or unmeasured.

### Cross disorder differential gene expression

Differential gene expression for each diagnosis group was calculated using a linear mixed effects model. Counts for all cell type within a brain region were derived from the Seurat counts matrix (Methods, Single nucleus RNA-seq Alignment and Filtering). To assess DGE by diagnosis within a given cluster, we subset the normalized counts to cells from that cluster to generate a per-cluster cell by gene counts matrix. Genes were then filtered using a minimum proportion, keeping only genes expressed in at least 10% of cells within any condition, calculated independently per cluster. We then calculate differential gene expression using a linear mixed effects model (LME) using lmerTest::lmer (Kuznetsova, 2017) with model formula ∼ clinical_dx + pmi + age + sex + number_umi + percent_mito + (1 | library_id). Resulting p-values were then FDR-adjusted across the number of genes measure in the DGE analysis. Depending on the experiment, different contrasts were used to measure DGE by disease. In some cases we contrasted one disease group, such as AD, with samples from all other diagnosis groups (such as PSP, bvFTD and control). In other cases, we generated one combined linear model to measure the effect on gene expression of each diagnosis group, contrasting each diagnosis with control (expression ∼ clinical_dx+ pmi + age + sex + number_umi + percent_mito + (1 | library_id)).

In several instances, as indicated in the text and figures (Figure 3K, Figure 4B,G,I, Figure S7C, Table S5) we used LME to measure differential gene expression between two specific clusters, within only samples of one disease samples (for example, comparing gene expression of INS_EX-2 vs INS_EX-5 in bvFTD cases, or between layer 2-4 vs layer 5-6 excitatory neurons). We subset the normalized counts matrix to only those only cells assigned to the cluster or cell subtypes and being compared (such as INS_EX-2 and INS_EX-5) and to only the libraries of the disease group of interest (such as bvFTD). As before, genes were then filtered using a minimum proportion, keeping only genes expressed in at least 10% of cells. The cells were subset to form two levels of a contrast labeled custom_split. We calculate differential gene expression on the counts matrix using a linear mixed effects model (LME) using lmerTest::lmer (Kuznetsova, 2017) with formula expression ∼ custom_split + pmi + age + sex + number_umi + percent_mito + (1 | library_id). Resulting p-values were then FDR-adjusted across the number of genes measure in the DGE analysis.

### Gene regulatory network analysis

We used a modified version of the SCENIC (Single-Cell rEgulatory Network Inference) approach (Aibar *et al*., 2017) for constructing GRNs from single-cell RNaseq data. Briefly, SCENIC contains three steps: (1) identify co-expression modules between TF and the potential target genes; (2) for each co-expression module, infer direct target genes based on those potential targets for which the motif of the corresponding TF is significantly enriched. Each regulon is then defined as a TF and its direct target genes; (3) the RAS in each single cell is calculated through the area under the recovery curve. We performed this analysis on four cell types (excitatory neurons, oligodendrocytes, astrocytes, microglia) from each brain region (INS, BA4, V1), using final cell clusters post-quality control, outlier removal, regression and normalization in Seurat (Butler *et al*., 2018). On disease and control cells combined, and for each cell type and brain region, we ran SCENIC. We first filtered genes to those expressed in at least 5% of cells. We then subsampled from each library to draw a similar number of cells from each library and disorder for the analysis. To achieve this, we sampled at the 15% decile range of counts across samples, and then performed stratified random sampling to keep this decile range of cells from each sample. Stratified random sampling starts off by dividing a population into groups with similar attributes. This method was used to ensure that different segments in a population are equally represented. We then applied the SCENIC package algorithm in R (https://github.com/aertslab/SCENIC) to generate gene regulatory networks and activity scores of every regulon in each cell type. To compare regulon activity across disorders, we used Shannon entropy algorithm described in (Suo et al., 2018) to quantify regulon specificity scores (RSS) for each regulon for each disease condition. Briefly, this analysis quantifies the differences between two probability distributions, scored from 0 to 1. The essential regulators are predicted to be those with the highest specific scores. This application has been used previously to define cell type specific regulons. Therefore, we first confirmed that our entropy-based scores reproduced known cell type specific TFs, by comparing microglia to all other cell types and observing top ranked microglia specific TFs to include *RUNX1* and *SP11* (Figure S6A-B), which are two known core microglial transcriptional regulators. We then modified this application to assign a specificity score for a regulon for each disorder (Figure 5-6).

### snATAC sequencing and data analysis

Nuclei were extracted as above simultaneously (BA4, INS) with 78 samples subjected to snRNA-seq balanced by age, PMI and cause of death (n = 8 – 11 per diagnosis and region pair as shown in Table S1). Libraries were aligned to reference genome with 10X Genomics software, Cell Ranger (cellranger-atac count; https://support.10xgenomics.com/single-cell-atac/software) then all libraries were aligned together using the cellranger-atac aggr function. We used ArchR version 1.0.2 (Granja JM*, Corces MR* et al. 2021) to filter out low quality cells through nucleosome banding score < 4, TSS < 2, minimum fragments < 1000, and blacklist region ratio > 0.1. We additionally dropped low-quality samples P1_7at1_7, I3_6_at, and I1_7 (Table S1), leaving 141038 AD, 136732 PSP, 145106 bvFTD and 152369 control high quality cells from each dataset.

We normalized with ArchR’s iterative Latent Semantic Indexing and Harmony batch correction on the sequencing and preparation batches. Clustering was performed using the Seurat clustering algorithm with resolution = 0.8 to generate 23 clusters. To annotate the clusters we first created a gene activity score matrix where gene expression levels are roughly computed from fragment counts within gene body elements. We then used the snATACseq gene activity matrix to integrate with our fully annotated snRNAseq clusters through Seurat’s ‘FindTransferAnchors’ function. We then used spearman correlation to assess how closely related the cluster-specific marker gene profiles from the snATACseq clusters (from the gene activity matrix) with our snRNAseq cluster marker gene profiles. This identified two snATAC-seq clusters corresponding to excitatory neurons (C2) and microglia (C7), which we used for further analysis.

### Peak calling and prediction of TF activity in snATAC-seq data

We merged reads from individual cells of snATAC clusters by disease condition and brain regions. Pseudo-bulk replicates were further created in ArchR using customized parameters to ensure balanced number of cells for all pseudo-replicates (ArchR::addGroupCoverage, minCell = 950, minRep = 4). Peak calling was performed using macs2 (https://github.com/macs3-project/MACS) in ArchR (Granja et al., 2021), resulting in a reproducible, non-overlapping peakset (ArchR::addReproduciblePeakSet). Differential accessible peak regions were determined by pair-wise comparisons of chromatin accessibility between disease and control cells. We identified significant TFs sourced from our SCENIC analysis and JASPAR via ArchR wrappers (addBgdPeaks, addDeviationsMatrix) for chromVAR (Schep AN, Wu B, Buenrostro JD and Greenleaf WJ, 2017), selecting TF motifs that were correlated with significant deviations in chromatin accessibility. To infer transcription factor activity, we performed TF footprinting analysis on peak regions differentially accessible in each disease using TOBIAS(Bentsen et al., 2020), as described in our previous study (Tian et al., 2022) (Wamsley et al., 2023 biorxiv) This method starts with Tn5 bias correction using the TOBIAS ATACorrect module, subtracting the background Tn5 insertion cuts highlighting the effect of protein binding. To match footprints to potential TF binding sites, and to estimate TF binding activity on its target loci, we applied TOBIAS BINDetect module to the corrected ATAC-seq signals within peaks, with TF motif PWMs used in TRASFAC Pro. Many transcription factors are represented by more than one motif. To avoid motif redundancy, we clustered the motifs based on their sequence similarity using TOBIAS ClusterMotifs, and chose one motif per TF that is most similar to others in the same cluster TOBIAS BINDetect compares the positions and activities of TF footprinted sites in disease or control per clusters. Each footprint site was assigned a log2FC (fold change) between two conditions, representing whether the binding site has larger/smaller TF footprint scores in comparison. To calculate statistics, a background distribution of footprint scores is built by randomly subsetting peak regions at ∼200bp intervals, and these scores were used to calculate a distribution of background log2FCs for each comparison of two conditions. The global distribution of log2FC’s per TF was compared to the background distributions to calculate a differential TF binding score, which represents differential TF activity between two conditions. A P-value is calculated by subsampling 100 log2FCs from the background and calculating the significance of the observed change. By comparing the observed log2FC distribution to the background log2FC, the effects of any global differences due to sequencing depth, noise etc. are controlled. To visualize, we used soGGI to plot TF footprints, and bar graphs to show the global TF footprint activity changes comparing disease versus control.

### Protein-Protein Interaction and Gene Ontology Analysis

To assess and visualize protein-protein interactions among module genes, we used STRING (version 11.5; (Szklarczyk et al., 2017)) with the following setting (organism: *Homo sapiens* for human data; meaning of network edges: confidence; active interaction sources: experiments and databases; minimal required interaction score: high confidence, max number of interactors to show: none; we reported either full STRING networks indicating edges representing both functional and physical protein associations, or physical STRING networks indicating only physical protein interactions based on databases, as indicated in Figures and Figures Legends). Protein-protein interaction enrichment p-values were reported as generated by STRING. Gene ontology enrichment represented within protein-protein interaction networks were prioritized and displayed along with their geneset enrichment FDR corrected p-values as generated by STRING, using whole genome as the default background. Data was exported and visualized using the Cytoscape software (Saito et al., 2012).

### Connectivity Map (CMAP) Analysis

For a given module, the top 100 module genes (ranked by T-statistic, LME) were used as input for the QUERY app in the Broad’s CMAP database, version CLUE (Subramanian *et al*., 2017). This signature was used to query 7,494 gene overexpression or knockdown experiments carried out across 9 cell lines for similar (positive connectivity score) or opposite (negative connectivity score) effects on gene expression signatures, incorporating Kolmogorov-Smirnov statistics (a nonparametric, rank-based pattern-matching strategy) as described(Subramanian *et al*., 2017). Per the CMAP website (https://clue.io), for each module-perturbagen pair, the connectivity score (*tau*) is a standardized percentile score that compares the similarity of the query geneset to the perturbagen compared to all other reference genesets in CMAP; such that 95 indicates that 5% of reference genesets show stronger connectivity to the perturbagen than the query dataset. For our analysis, we used the mean “connectivity scores” which is calculated from the combining data generated independently in 9 cell lines.

### AD GWAS risk variant enrichment

Summary statistics for genome-wide association studies for AD (Lambert *et al*., 2013) were used as an input for MAGMA (v1.08bb) (de Leeuw et al., 2015) for gene annotation to map SNPs onto genes (with annotation window = 20) and the competitive gene set analysis was performed to test module associations with GWAS variants (permutations = 100,000). Genesets analyzed included marker genes of differentially disease enriched clusters (LME, T-statistic>2); as indicated in the text or accompanying figure legends. FDR correction was applied across competitive p-value outputs from MAGMA for all genesets tested.

### Module enrichment

We implemented a Fischer’s exact test for geneset enrichment analysis. We compared cluster-specific and DGE by disorder (Log2FC > 0.2 in one disease vs all others, LME, see above section) with genes assigned to TF regulons by SCENIC (see Table S6). To compare cluster-specific marker gene expression (T-statistic >2, LME, see above section) to published disease-associated cell types and markers, we used published genesets and markers shown in Table S3.

Unless otherwise stated all statistical test were run using R (4.1.0). P-values are reported with post-hoc FDR for multiple testing correction.

## Supplementary Information

**Supplementary Tables 1: Sample metadata**

**Supplementary Tables 2: Cluster Marker Genes**

**Supplementary Table 3: Differential Composition**

**Supplementary Table 4: Comparison with Published Clusters**

**Supplementary Table 5: Cluster-specific Differential Gene Expression**

**Supplementary Table 6: Gene Regulatory Networks**

**Supplementary Table 7: Chromatin Analysis**

**Supplementary Figure 1:** (**A**) Boxplot of sample characteristics measured across nuclei used in final analysis separated by brain region and disease condition, including subject data (post-mortem interval (PMI), age), and data measured per single nuclear library (number of unique molecular identifiers (UMI), percent reads mapped to mitochondrial genome (mito-DNA); mean +/- quantiles). (**B**) Box plot showing percent of each of the 9 most prominent cell types averaged over each sample (mean +/- quantiles). (**C**) UMAP based on reference-based mapping of motor cortex samples (BA4, all conditions, Azimuth (see Methods), Human Motor Cortex as reference (BICCN)) demonstrating 100% of reference cell subtypes ((Bakken *et al*., 2021)). (**D**) UMAP showing results of unsupervised clustering of excitatory neurons from one brain region (BA4) and all disease conditions (LIGER subclusters; Methods). (**E**) Sankey plots showing assignment of clusters to matched reference cell classes, and, for excitatory neurons, inhibitory neurons and astrocytes, to reference cell subclasses (Azimuth (see Methods), Human Motor Cortex as reference (BICCN)).

**Supplementary Figure 2:** (**A-G**) Hierarchical clustering for 7 major cell types, showing grouping of related clusters from three brain regions based on marker gene overlap within cell type (Methods, Table S2). As shown in legend, clusters are colored and labeled based on their brain region of origin (V1 white, BA4 grey, INS black; as shown in key), and their numeric cluster identifier (matching unique cluster identifier “brain region_ celltype_ numeric”), for (**A**) IN = inhibitory neurons, (**B**) OL = oligodendrocytes, (**C**) AST = astrocytes, (**D**) OPC = oligoprogenitor cells, (**E**) END = endothelial cells, (**F**) Pericytes, and (**G-H**) MIC = microglia. IN, OL and AST subclasses are labeled based on reference-based classifiers (Methods; for example OLIGO L3-L6 *ENPP6* (Bakken *et al*., 2021), Table S2). For each cell type, we include a heatmap indicating cluster-specific differential expression of a selection of reference-based classifier genes and genes that distinguish related clusters (log2FC vs other clusters from same brain region and cell type, LME; Table S2). Representative marker genes that distinguish cluster groups are also shown and used to label groups of related clusters for each cell type (see Table S3 for markers listed with references). For example *ENPP6,* a marker of putative newly formed OL (Xiao *et al*., 2016), *RBFOX1*, *PLP1* indicating myelinating OL, and *BCAS1* marking early pre-myelinating and/or disease associated OL (Fard *et al*., 2017; Hughes and Stockton, 2021; Xiao *et al*., 2016) (See Table S3 and S2 for additional markers). Clusters significantly depleted (-) or enriched (+) in disease cells are indicated with a colored line (AD red; bvFTD blue, PSP gold; all disorders black) based on strict criteria (LimmaVoom (Methods); see Table S3 for differential cluster composition summaries and complete results). For example, two OL clusters in B are upregulated in bvFTD specifically while another is upregulated in PSP specifically, one MIC cluster is upregulated in AD specifically, and one AST cluster is upregulated in bvFTD specifically. Statistical criteria for disorder-distinct results shown are abs(beta) > 0.75 and FDR<0.1 for one diagnosis vs all other conditions (no star), or FDR <0.1 in only one diagnosis vs control (indicated by *). For selected trends reported in Table S3 that do not meet these strict significance thresholds, FDR corrected p-values are shown. Statistical criteria for shared disease-associated findings shown are abs(beta) > 1 for each disorder vs control, and FDR <0.05 for all disorders combined vs controls. FDR corrections were applied across clusters of the same cell type and brain region (see Table S3). For (**H**) the cluster DGE results compare markers previously associated with homeostatic microglia (Chen and Colonna, 2021) and related markers, demonstrating broad expression of some markers across multiple clusters (*CSF1R*, *P2YR12*), and more discrete expression of *SLCO2B1*, *EIF4G3*, *TGFBR1* in microglia depleted in disease (INS_MIC-0).

**Supplementary Figure 3: Reproducible and shared disease associated cell types in AD compared to bvFTD and PSP.** (**A**) Reproducible depletion of *RORB*+ *NEFM*+ layer 4/5 excitatory neurons is specific to AD samples and not observed in PSP or bvFTD in BA4, and in control samples, neurons selectively depleted in AD (BA4_EX-4) show higher expression of *RORB* compared to layer 4/5 neurons that share subclass specific marker genes but are not depleted in AD (heatmap showing differential expression of layer specific marker genes; bar plot showing differential composition of BA4_EX-4 neurons in AD, bvFTD and PSP samples vs control (LME) and bar plot showing differential expression of *RORB* in BA4_EX-4 neurons compared to BA4_EX-7 neurons in control and AD samples (***FDR<9e-10 corrected for 26,084 transcripts). (**B**) INS_IN-10 are of the same subtype as AD-affected interneurons reported in prior work (Mathys *et al*., 2019) based on high differential expression of cluster-specific marker genes compared to other INS interneurons (LME), and they are differentially enriched in AD samples more than PSP, bvFTD and controls (limma *p-value =0.48, Table S3). (**C**) Astrocytes in total robustly downregulate *SCL1A3* in all disease conditions and brain regions, similar to what has been previously reported in AD (Leng *et al*., 2021); LME T-statistic shown, Methods). (**D**) Characterization of a shared disease associated excitatory neuronal clusters from V1 based on differential composition analysis (limma, ***FDR <0.001 with 12 comparisons (V1_EX clusters), Table S3), differential expression of layer specific marker genes (Methods), direct PPI plot showing enriched gene ontology and PPI enrichment p-value (STRING, showing enrichment FDR corrected p-values compared to whole genome), and overlap with gene expression signatures of excitatory neurons present in iPSC-derived organoids generated from V337M patient lines (Bowles *et al*., 2021)(Fisher’s test, ***FDR<0.001 over 6 comparisons shown). (**E**) Characterization of shared disease-associated genes in *MEIS2* interneurons (INS_IN0,6) including differential expression of marker genes (LME, normalized expression t-statistic, Table S2), differential composition across disorders compared to controls (limma, *FDR<0.1 corrected for comparison across 11 INS_IN clusters), and genes up- and down-regulated in INS_IN-0 samples in all disorders (below, showing Log2FC vs controls per diagnosis group, *FDR <0.05 corrected over 10,772 genes (INS_IN-0) and 10,432 genes (INS_IN-6), Table S5), and their functional PPI plots showing enriched gene ontology and PPI enrichment p-value (STRING, showing enrichment FDR corrected p-values compared to whole genome). (**F**) Characterization of shared disease-associated microglia including differential composition by disorder (limma; top, * indicates FDR < 0.05 for differential composition in all diseases vs control, corrected over 10 INS MIC clusters) differential expression of marker genes (Log2FC vs other clusters from same brain region and cell type, LME; Table S2), (**G**) overlap of cluster-specific markers with published disease-associated microglia clusters defined in AD (Mathys *et al*., 2019) and MS (Schirmer *et al*., 2019) (Fisher’s exact test, FDR over 20 comparisons *<0.05, **<0.01, ***<0.001), (**H**) overlap of cluster-specific marker genes with functional gene perturbation class signatures (CMAP connectivity score (Subramanian *et al*., 2017), Methods), showing INS_MIC-3 specifically inversely correlated to WNT and PI3K pathway loss-of-function (LOF) and (**I**) disease-associated changes in gene expression shared across disorders in INS_MIC-3 showing up-regulation of WNT and PI3K pathways (top 100 genes ranked by t-statistic, LME, Table S5) and functional PPI plots of significantly up-regulated genes in disease (Log2FC > 0.2 and *FDR < 0.05 measured for each disease vs control, corrected over 8,220 genes) showing their enriched gene ontology and PPI enrichment p-value (STRING, showing enrichment FDR corrected p-values compared to whole genome).

**Supplementary Figure 4** (**A**) Boxplot showing proportion of cells in a given cluster per cell type and brain region, per sample library, for clusters with differential compositions in disease cases (boxplot shows mean +/- quartile). (**B**) Marker gene expression distinguishing microglia clusters from BA4 (Log2FC vs other clusters from same brain region and cell type, LME; Table S2; *FDR<0.05, **FDR<0.01, *** FDR<0.001, corrected over 1942 genes, Table S2). (**C**) Overlap of cluster-specific marker genes with functional gene perturbation class signatures (CMAP connectivity score, Methods). (**D**) Overlap of cluster-specific markers compared to published disease-associated microglia clusters defined in microglia isolated from fresh brain tissue (Olah *et al*., 2020) and AD prefrontal cortex (Mathys *et al*., 2019); Fisher’s test, left 27 comparisons, right 12 comparisons, as shown). **(E)** Correlation between percent BA4_MIC-4 in disease samples (vs total BA4 microglia counts) per library compared to sample qualitative tau neuropathology score (n = 15, samples colored by diagnosis; controls excluded from analysis). (**F**) Functional PPI plots showing enriched gene ontology and PPI enrichment p-values of genes upregulated in BA4_MIC-4 in bvFTD samples (cross disorder LME, top 150 genes ranked by t-statistic, Table S5; STRING FDR corrected p-values compared to whole genome). (**G**) Differential composition of BA4 microglia clusters in bvFTD samples compared to all other conditions (limma, *FDR=0.025 corrected over 4 clusters). (**H**) Differential gene expression in BA4_MIC-1 in bvFTD vs other conditions as indicated (LME, FDR corrected for 2,612 genes detected). (**I**) Correlation between percent BA4_MIC-1 in disease samples (vs total BA4 microglia counts) per library compared to sample qualitative tau neuropathology score (n = 10, controls applied over number of 5 comparisons). Regulons are ordered from left to right based on specificity score rank (Table S6), with highest ranking (most disease associated) TF regulons list to left. RSS ranks are as follows: BA4_AST for bvFTD (*SOX5* (1), *SOX12* (3), *NFAT5* (5), *SREBF1* (38), *TCF7L2* (40)), V1_AST for PSP (*ZMAT4* (1), *NR3C1* (3), *CUX2* (5), *FOXP1* (38), *JUN* (40)), INS_OL for bvFTD (*RELA* (1), *NEFLE* (2), *ELK4* (3), *FOXN3* (57), *STAT1* (71)), BA4_OL for PSP (*ATF4* (1), *NFE2L1* (2), *OLIG2* (3), *SOX10* (4), *STAT3* (60)), BA4_MIC for bvFTD (*IRF8* (1), *RUNX1* (2), *NRC31*(9), *IKZF1* (21), *SPI1* (24)). (**D**) TF regulon plot showing regulon target genes (edge length proportion to (1-GRN score) that are up-regulated (log2FC) in clusters in which they are enriched. Here nodes are colored scaled to their disorder-specific differential gene expression (log2FC, LME, Table S5). (**E**) Overlap of genes in *SPI1* regulons comparing the regulon from V1 (blue) and BA4 (red) microglia. SPI1 regulon genes that overlap between regions are listed at the intersection, and those that are unique to each brain region are shown as part of functional PPI networks. Enriched gene ontology pathways are colored as indicated in the legend. The legend also shows enrichment p-values listed to the right of each GO term (STRING FDR corrected p-values compared to whole genome).

## Data Availability

Data will be deposited for public use at publication. Data will be made available for review.

## Code Availability

https://github.com/rexachgroup/snseq_nd_liger/

