## Supplementary material for "Disease-specific selective vulnerability and neuroimmune pathways in dementia revealed by single cell genomics": Supp Figures

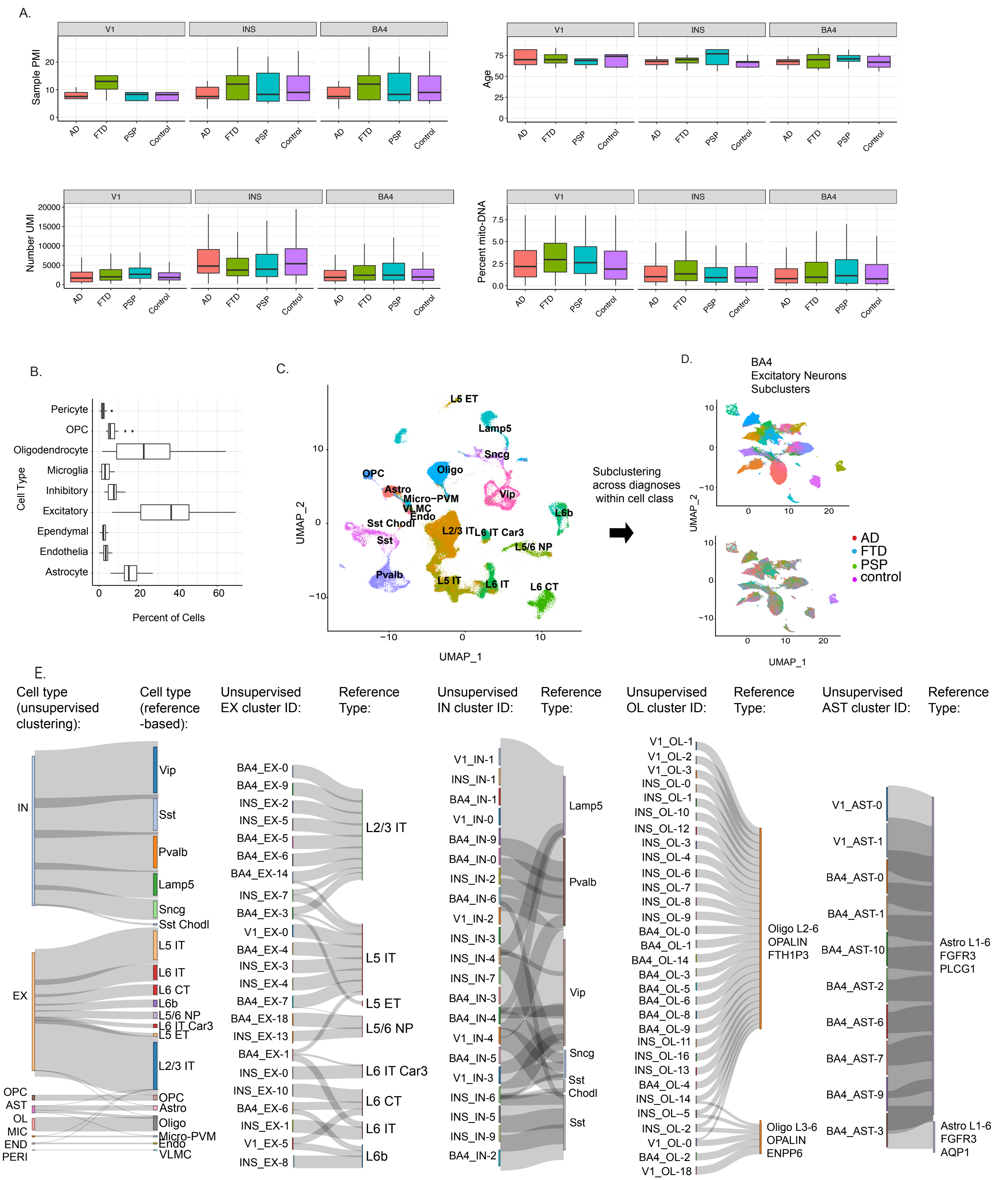


Supplementary Figure 1

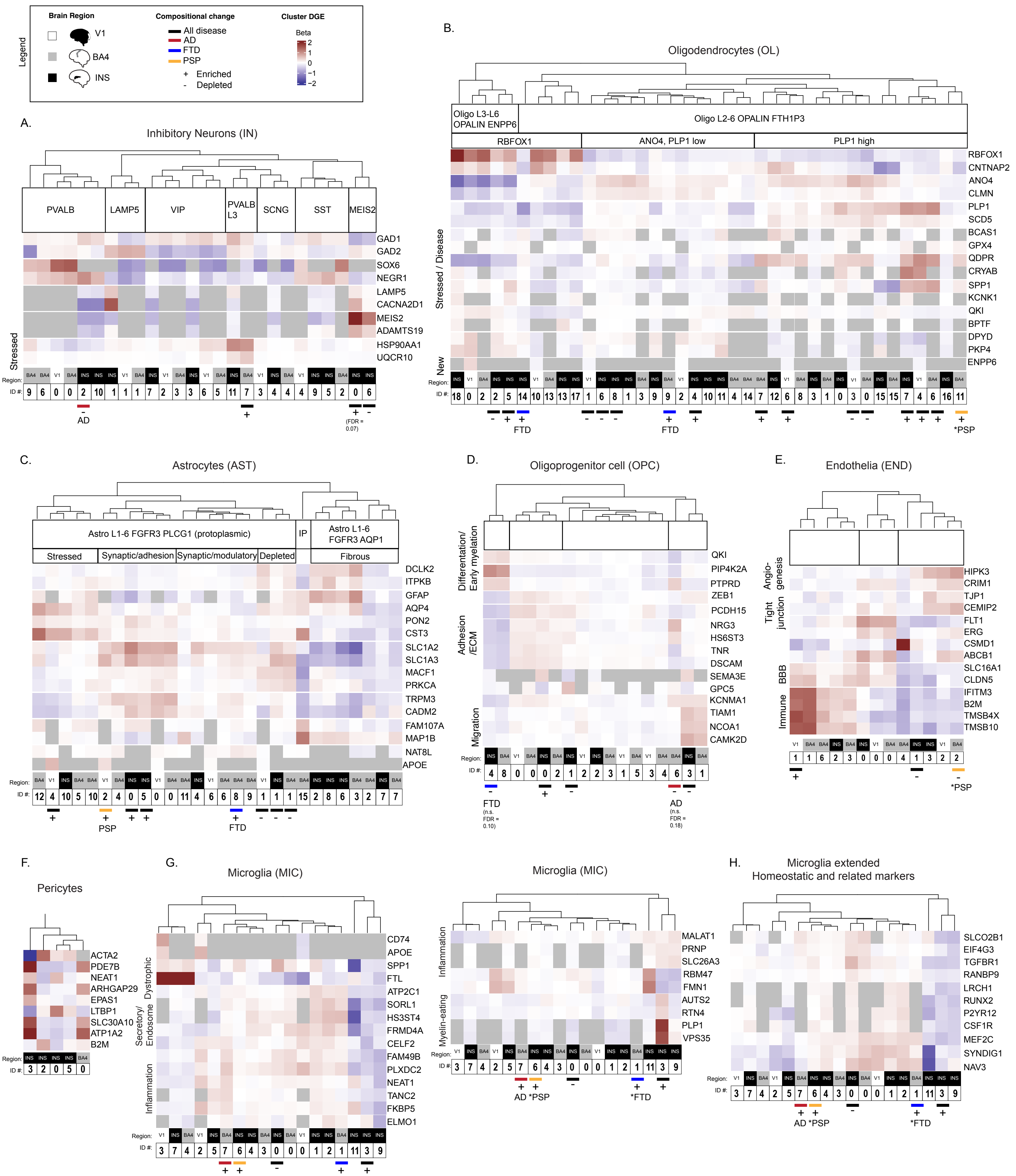


Supplementary Figure 2

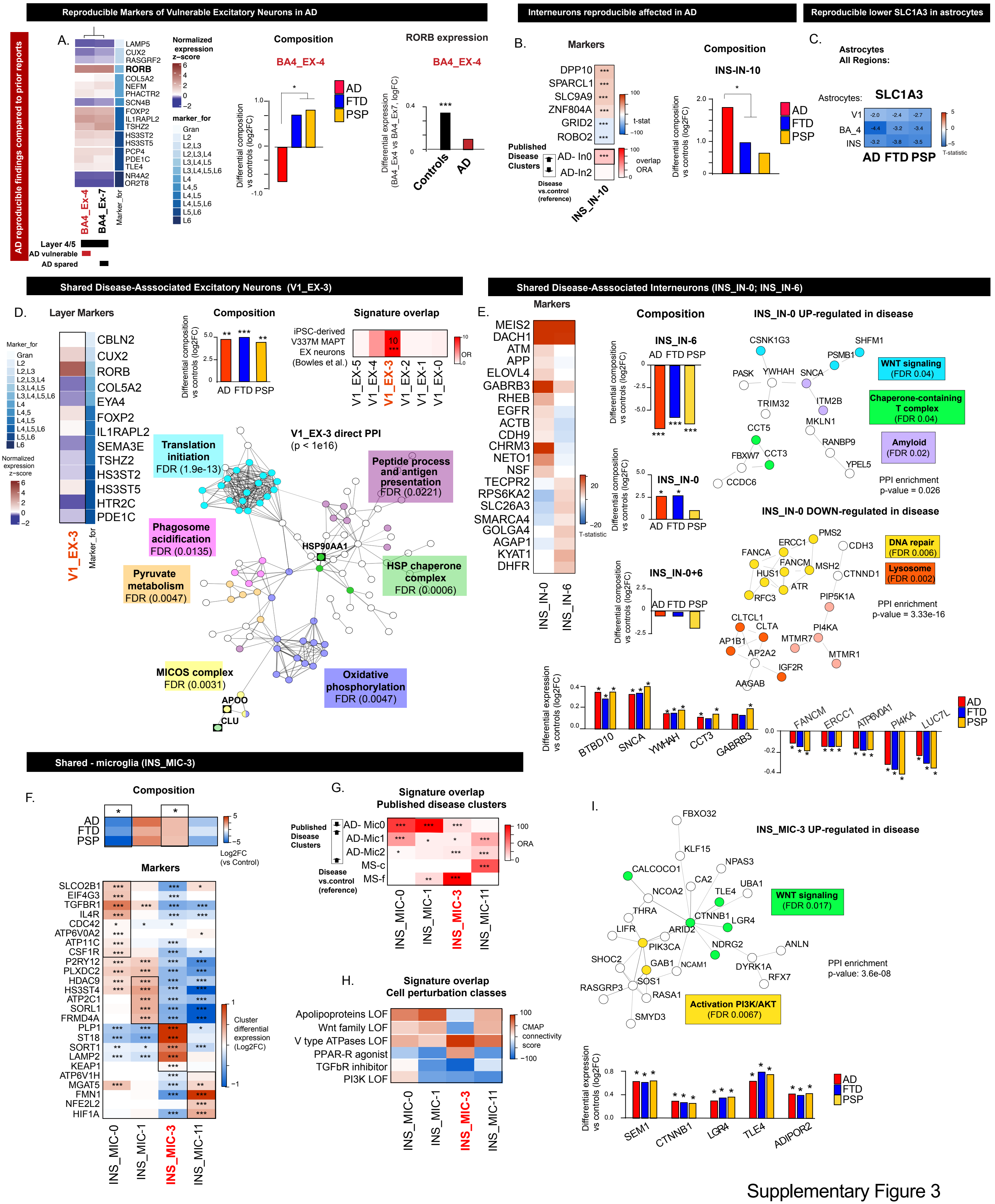


Supplementary Figure 3

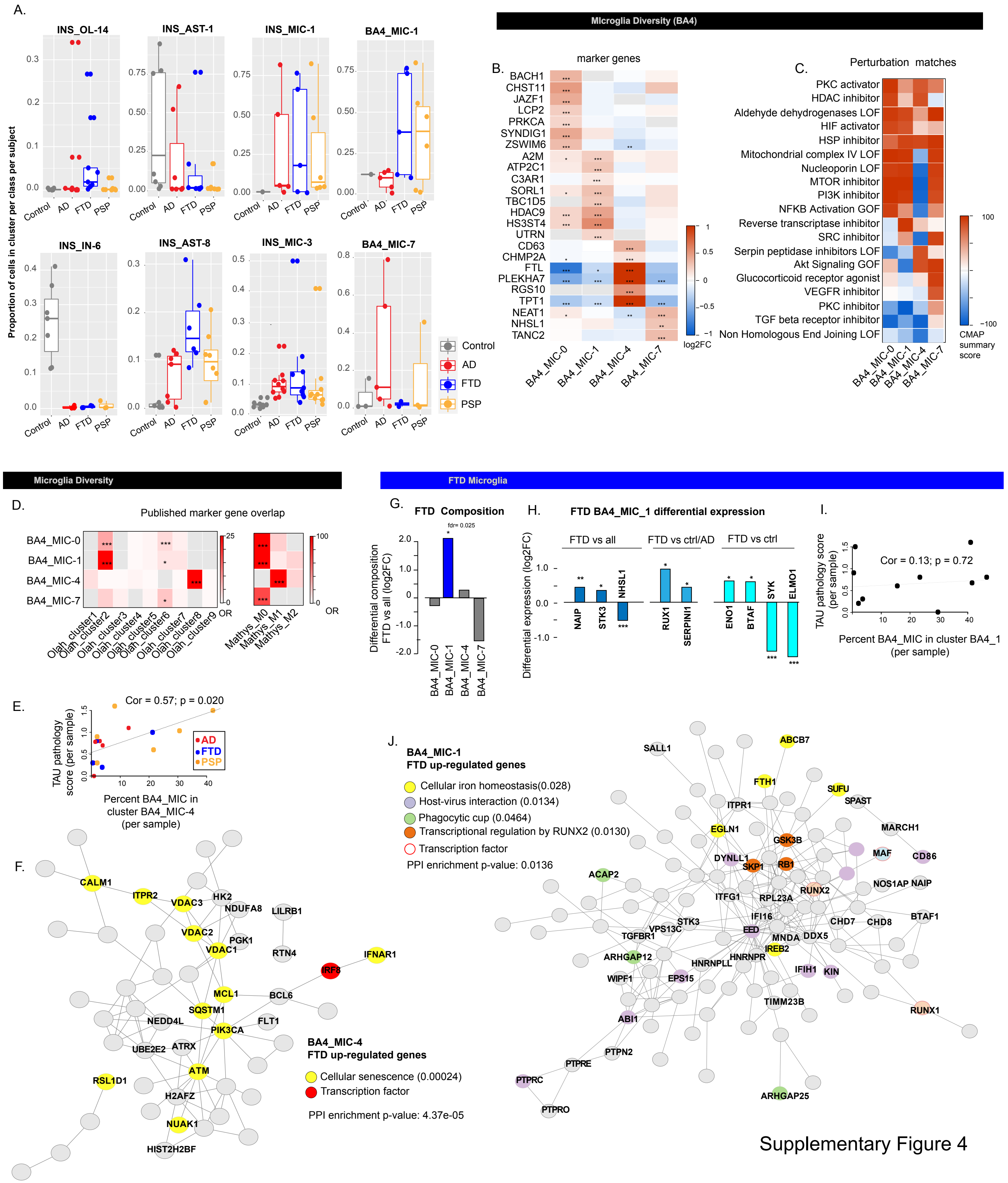


Supplementary Figure 4

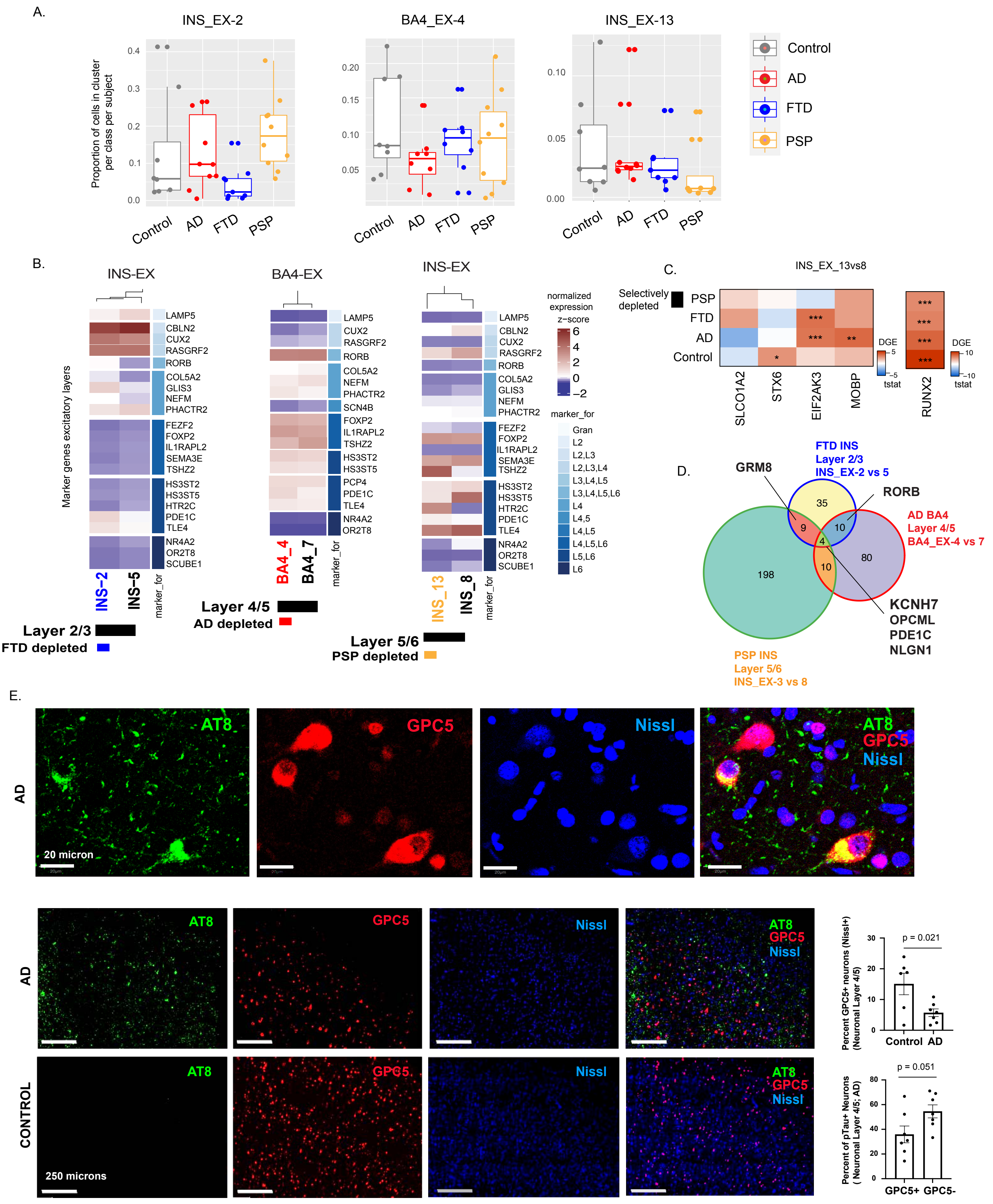


Supplementary Figure 5

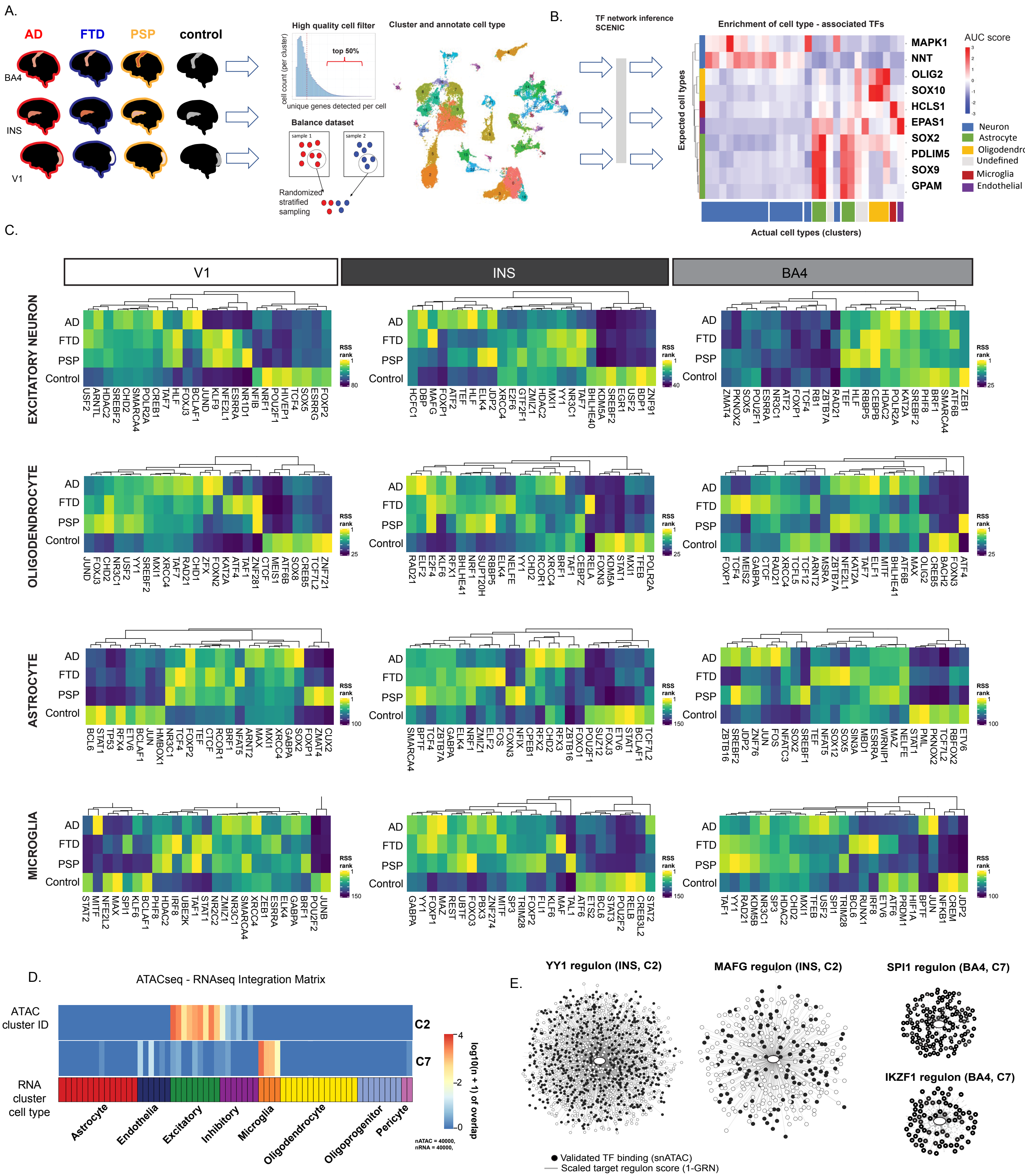


Supplementary Figure 6

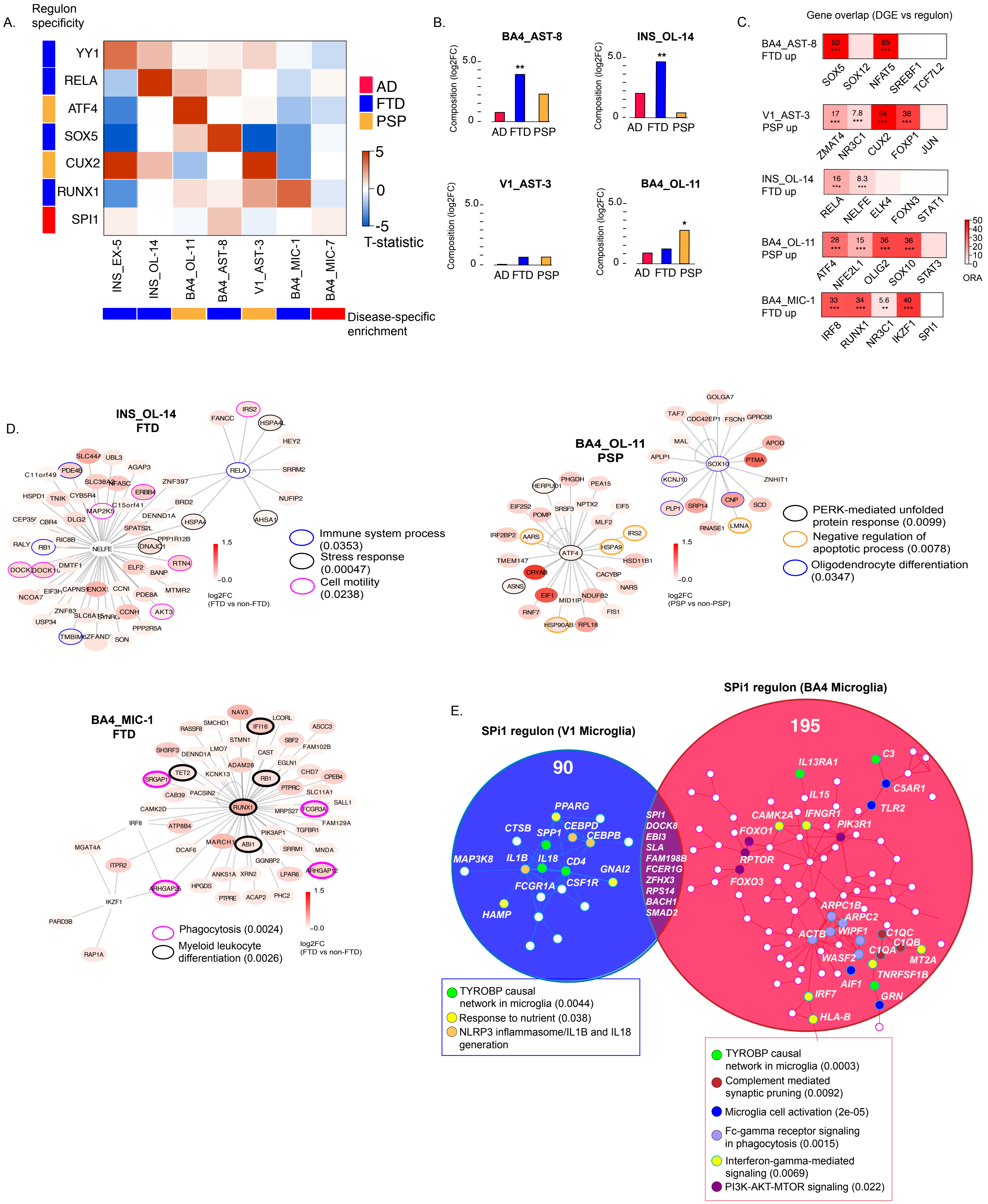


Supplementary Figure 7
